# Dynamic gene-body chromatin accessibility is associated with a stage-dependent transcriptional upregulation during germination of fission yeast spores

**DOI:** 10.64898/2026.11.24.769058

**Authors:** Yuki Shogaki, Hayato Tsuyuzaki, Ryutaro Aonuma, Chikako Sakanashi, Masahito Hosokawa, Haruko Takeyama, Yasutaka Kakui, Masamitsu Sato

## Abstract

Variations in the expression of histone genes influence the process of breaking dormancy in fission yeast spore cells. Since histones play a crucial role in chromatin organization, alterations in their expression impact chromatin states during spore germination, where such changes are vital. Although chromatin accessibility state has been studied in proliferating cells, there has been no specific research on spores. In this study, using ATAC-seq to examine chromatin accessibility in fission yeast spores, we found that ATAC-seq peaks frequently appeared as broad regions in gene-body in addition to promoter-linked narrow regions. Narrow peaks representing high chromatin accessibility at transcription start sites (TSSs) observed in proliferating cells were maintained in dormant spores as well as germinating spores, whereas broad peaks that cover gene-body were redistributed among different genes as germination progressed. The transition of broad open chromatin on gene-body was often linked with a stage-dependent transcriptional upregulation determined by scRNA-seq throughout dormancy breaking. Thus, the spore-specific chromatin alterations contribute to transcriptional changes observed during germination.

## Introduction

Eukaryotic cells proliferate through cycles of cell division. However, not all cells continuously proliferate; in unsuitable environments such as nutrient starvation, they may enter a dormant state^1^. Dormant cells can resume proliferation when placed in an environment conducive to growth. In yeast, dormancy takes two forms: quiescent cells arrested in G0 and spores. When proliferating cells encounter nutrient starvation, they arrest the cell cycle in G1 phase and then enter a quiescent G0 state^2,3^. Alternatively, starved diploid cells, or haploid cells mated with cells of the opposite mating types, undergo meiosis and generate spores as a dormant form^4^.

Spore germination is the process to resume cell growth from dormant spores, starting from dormancy breaking in response to glucose^5,6^. Because individual fission yeast spores break dormancy at different timing after nutritional feeding, we previously conducted transcriptome analysis of single spores using single-cell RNA-seq (scRNA-seq) technique and prepared a virtual germination timeline^7,8^. This analysis showed that expression of *hht1/h3.c1*, one of the genes encoding histone H3, temporarily decreases during dormancy breaking and recovers later, and that the reduction of Hht1/H3.c1 in histone H3 facilitates the germination process. However, the mechanism by which the reduction of Hht1/H3.c1 contributes to germination through chromatin state remains unexplored.

Dormancy is accompanied by changes in chromatin. In carbon-starved quiescent budding yeast, genome-wide chromatin compaction occurs^9,10^. Nitrogen-starved G0 fission yeast cells have small nuclei with the repression of transcription and mild histone deacetylation^11^. Nucleosomes, in which genomic DNA is wrapped around histone octamers, serve as fundamental units of chromatin^12^, and chromatin structure is implicated in the regulation of gene expression^13^; therefore, histones are posited to play a role not only in the compact storage of chromosomal DNA within the cell but also in the regulation of gene expression. Transcriptionally active genes and regulatory regions are known to adopt an accessible state, referred to as open chromatin^14^. Based on these understandings, we hypothesized that quantitative fluctuations in Hht1/H3.c1 levels during germination would affect gene expression profiles through alterations in chromatin accessibility.

To validate this hypothesis, the chromatin configuration upon germination needs to be examined. Chromatin states have been assessed by enzymatic digestion of genomic DNA with DNase or MNase, followed by molecular cloning or microarray^15,16^, and by DNase-seq, MNase-seq, FAIRE-seq, and ATAC-seq in a genome-wide manner across diverse species^17–20^. Among these techniques, we chose ATAC-seq (Assay for Transposase-Accessible Chromatin using sequencing), which is characterized by its simplicity and ability to provide both chromatin accessibility and nucleosome positioning in cells and tissue sections from various species, including fission yeast vegetative cells^20–22^. To our knowledge, however, ATAC-seq has not been applied to yeast spores due to the rigid spore wall, which limits the access of transposase Tn5 to chromatin. In our previous study, the expression of spore-specific ß-1,3-glucan synthase Bgs2 was reduced to make the spore wall fragile^7,23,24^. The *bgs2* mutant spores readily burst in a hypotonic buffer. We therefore refined the ATAC-seq protocols for spores using the the *bgs2* mutant.

Here, we monitored the dynamics of histones during germination using GFP-tagged proteins, and determined the profiles of chromatin accessibility in spores at 0, 1, and 3h after germination induced by nutritional feeding in comparison with that in proliferating vegetative cells. We found that multiple histone subunits transiently decreased as observed for Hht1/H3.c1. Our ATAC-seq showed that the transcription start site (TSS)-proximal nucleosome depleted regions (NDRs) were maintained throughout germination, whereas increased chromatin accessibility in gene bodies (coding sequences) differed among the stages of germination. These genes associated with increased chromatin accessibility overlapped with the genes upregulated in a stage-dependent manner in our previous scRNA-seq data. These results shed light on changes in chromatin accessibility associated with gene expression during spore germination.

## Results

### Histone-GFP temporally fluctuates for major components of the histone octamer

In earlier studies, it was found that among the three histone H3 genes in fission yeast (*hht1*, *hht2*, *hht3*), only *hht1* (histone H3.c1) showed a temporary reduction in mRNA and protein levels, while *hht2* and *hht3* remained unchanged^7^. We aimed to investigate whether this transient reduction in mRNA/protein level during germination is unique to *hht1* or if it is a common characteristic also seen in other histones besides H3. Then, protein levels of histone H2A, H2B and H4 during germination were assessed using *S. pombe* strains in which the GFP gene was fused with histone genes at their endogenous loci (**Fig. 1a**). Hta1(H2A)-GFP displayed a sharp decline in GFP fluorescence intensity, as Hht1(H3.c1)-GFP did, from the start of germination induced by nutritional feeding until 5 hours, after which the Hta1-GFP signal regained. The fluorescence of Htb1(H2B)-GFP and Hhf1(H4)-GFP did not decrease as markedly as that of Hta1-GFP; however, they similarly exhibited a temporary reduction during the middle of the germination process. Furthermore, as the brightness of Hht1-GFP increased during germination, the area exhibiting GFP fluorescence also expanded. If the area of histone GFP fluorescence expanded without a corresponding rise in expression levels, the GFP brightness would diminish. However, the concurrent increase in both area and brightness suggests an elevation in histone GFP expression levels. We assessed the relative temporal changes in GFP fluorescence intensity and the area where GFP fluorescence was detected for these observations. (**Figs. 1b and S1**). It was consequently observed that, although not as prominent as H3, other histones also exhibited a temporary decrease in fluorescence intensity during the initial stages of germination, followed by an increase in the later stages. Similarly, the region showing GFP fluorescence displayed an expansion in fluorescence coverage alongside an increase in fluorescence intensity for all other histones. It is suggested that the decrease in various histones, including but not limited to histone H3, might affect the chromatin structure of spores. This prompted us to investigate the impact on chromatin configuration during germination through ATAC-seq on spores.

**Figure 1.**
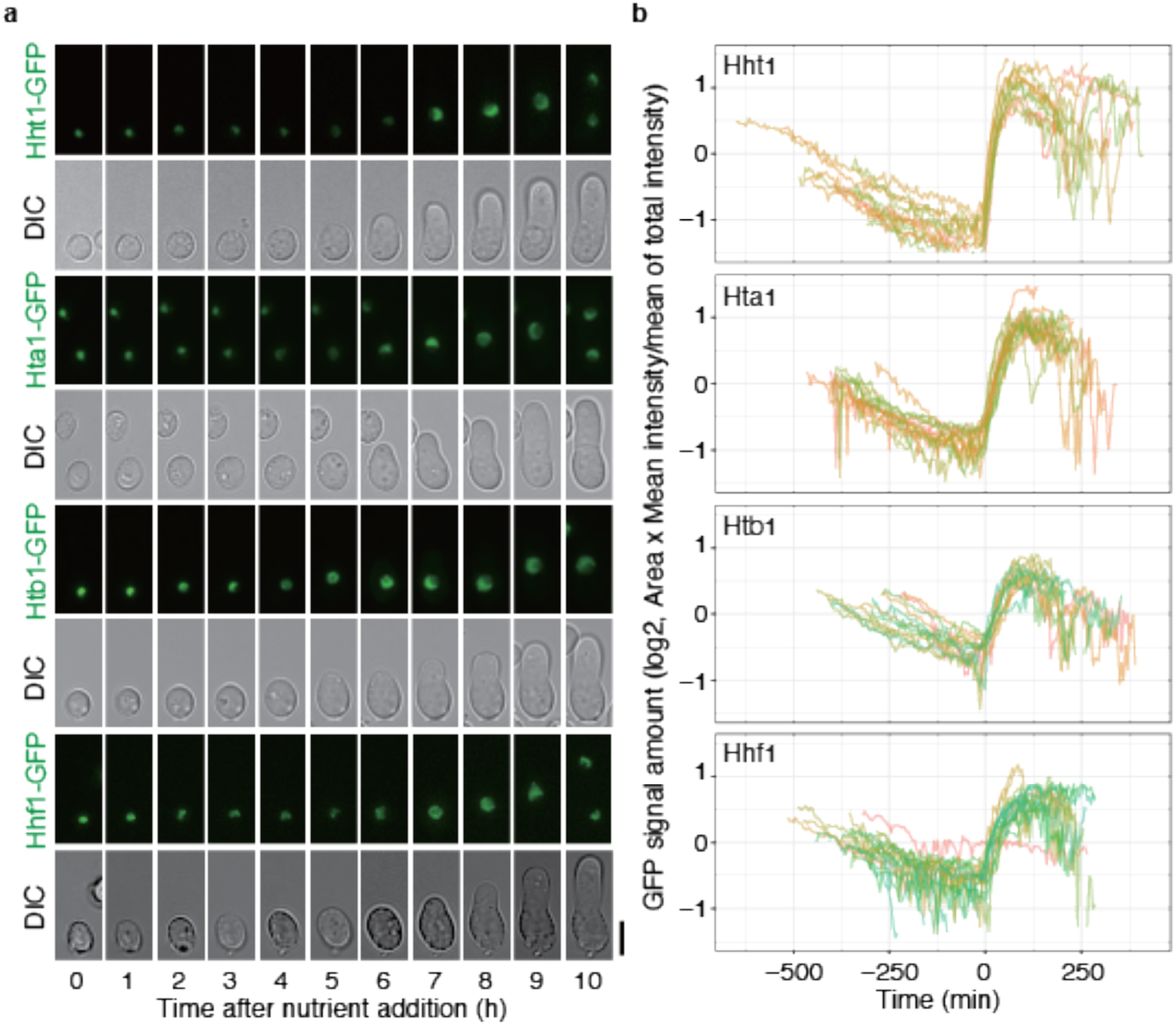
The majority of histone genes demonstrate variability in expression levels during the germination process. **(a)** Behavior of histone proteins tagged with GFP during the process of germination. Live-cell imaging was conducted for a duration of 10 h following the initiation of germination, which was induced by nutrient supplementation. Scale bar = 5 µm. **(b)** The temporal variation in GFP fluorescence intensity was monitored during the observation period. Time 0 (min) is designated as the point at which a discernible change in brightness was observed. n = 15 cells (Hta1-GFP), 14 (Htb1-GFP), 15 (Hht1-GFP) and 18 (Hhf1-GFP).

### Optimization of ATAC-seq protocols for *S. pombe* spores

ATAC-seq has been previously adapted for use in *S. pombe* vegetative cells^21,25^; however, its application to spores has not been reported. In this study, we refined the ATAC-seq protocols (see **Methods**; **fig. S2a**) by utilising the *bgs2* mutant, which had been previously employed in our spore scRNA-seq research^7^. Due to the fragile spore walls of the *bgs2* mutant, suspending the spores in Tn5 reaction solutions resulted in enzymatic interaction with the genomic DNA. Consequently, tagmented genomic DNA fragments were amplified as efficiently as those from spheroplasts. Two biological replicates were conducted for four distinct conditions: vegetative cells (spheroplast, see methods for details), dormant spores, and spores induced to germinate for either 1 hour or 3 hours (referred to 0h, 1h and 3h hereafter). The determination of the number of PCR cycles during library preparation, aimed at approximately equalising the sizes of libraries, was conducted following the methodology outlined by the previous study^20^. The specific number of additional PCR cycles incorporated during the library preparation for each sample is documented in **Supplementary Table 1**. The percentages of duplicates resulting from PCR amplification ranged from 0.28 to 0.67 (Duplication rate, **,Table S1**). This range is comparable to the previously reported range of 0.19 to 0.54, indicating that the duplication rates do not present significant concerns. The library size obtained for one sample was 1.81 million reads, whereas the sizes for other samples ranged from 3 to 8 million reads. Given that previous studies reported library sizes ranging from 3 to 11 million^21,25^, we ascertained that the library sizes achieved through our method fall within an acceptable range.

### Chromatin accessibility profiling of fission yeast spores

Upon analyzing the distribution of sequenced read sizes, it was observed that fragments measuring 200 base pairs (bp) or less constituted the majority (**fig. S2b**), indicating that reads derived from multiple nucleosomes are either present in low proportions or virtually absent. Fragments corresponding to sizes of mono-nucleosome (130-200 bp) and nucleosome-depleted region (NDR, 38-100 bp) were selected for the downstream analyses. To assess the similarity between libraries, the correlation of read counts mapped to 1 kilobase (kb) bins was calculated (**fig. S2c**). A high correlation was observed between the two biological replicates for each condition (Spearman’s ρ: 0.80–0.88). Furthermore, our vegetative conditions were highly correlated with the published datasets^21,25^, validating our datasets. A low correlation (Spearman’s ρ: 0.38–0.59) was noted between vegetative cells and spores, while the correlation coefficient was high (Spearman’s ρ: 0.74–0.88) among dormant (0h) and germinating spores (1h and 3h). These results suggest differences in chromatin configurations between these conditions. To elucidate the differences between spores and vegetative cells, the proportion of sequence reads mapped to each chromosome was assessed. (**Fig. 2a, Table S2**). If the sequencing reads are uniformly distributed across the genome, the quantity of mapped reads on each chromosome should be proportional to the chromosome length; thus, longer chromosomes are anticipated to exhibit a greater number of mapped reads. In vegetative cells, more reads were mapped to longer chromosomes in principle –– chromosome I, 30%; chromosome II, 25%; and chromosome III excluding the ribosomal DNA (rDNA) region, 15%. As the three chromosomes of fission yeast measure approximately 5.7 megabases (Mb) for chromosome I, 4.6 Mb for chromosome II, and 2.5 Mb for chromosome III, these proportions are comparable to the expected proportions from their lengths. Notably, approximately 30% of the reads were specifically mapped to the rDNA region located at the termini of chromosome III, which was the most abundant comparable to the mapped rate onto the entire chromosome I, the longest chromosome (Veg, **Fig. 2a**).

**Figure 2.**
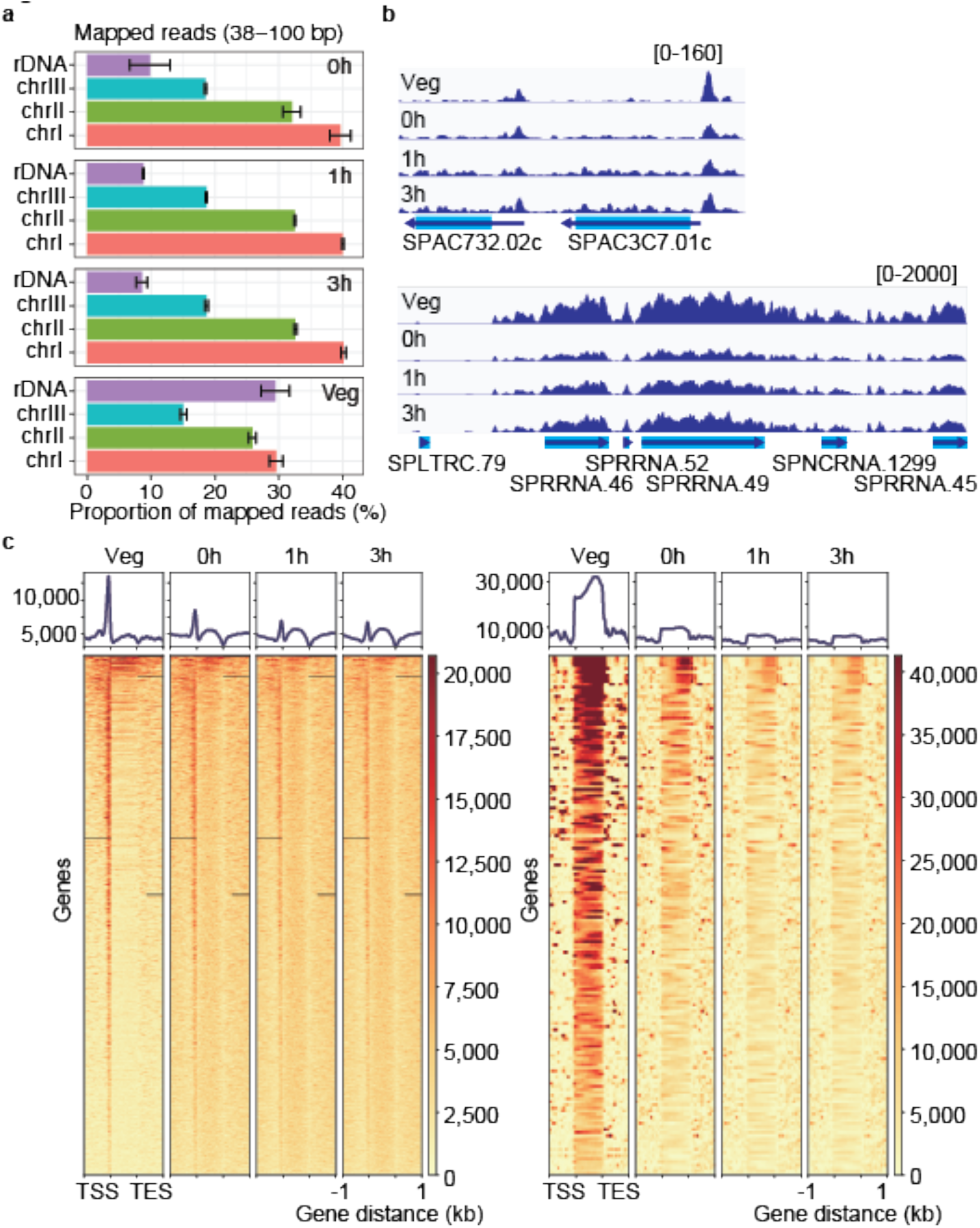
ATAC-seq reads in spores were predominantly enriched in coding sequence regions. **(a)** The proportion of reads mapped to each chromosomal region was calculated, with read sizes ranging from 38 to 100 bp. Values for chromosome III do not include reads mapped onto the rDNA region. Error bars represent two biological replicates. **(b)** An illustrative example of the distribution map of ATAC-seq signals aligned to the reference genome. A representative gene region, depicting two coding genes (SPAC732.02c and SPAC3C7.01c) transcribed in the reverse direction (top). A typical region containing clustered rDNA genes (SPRRNAs, bottom). Locations of coding sequences and directions of transcription are shown as boxes and arrows, respectively. **(c)** The vertical arrangement of read distribution per gene spans from the transcription start site (TSS) to the transcription termination site (TTS). Each gene was depicted with its length adjusted to position its transcriptional region between TSS and TTS. Distribution of all genes without non-coding RNAs (left), and the distribution of tRNA genes exclusively (right). As the genes in each category are arranged vertically, the number of genes displayed in the left and right panels differs. n = 5478 for CDS, n = 196 for tRNA.

In spores, by contrast, percentages of mapped reads onto each chromosome increased (chromosome I, 40%; chromosome II, 30%; and chromosome III excluding rDNA, 20%) compared to vegetative cells (0h, 1h and 3h, **Fig 2a**). Instead, percentages mapped to the rDNA region decreased to approximately 10%. After excluding mapped reads on rDNA regions, the distributions of mapped reads among the three chromosomes were proportional to their lengths. Therefore, a distinguishing characteristic of chromatin structure between vegetative cells and spores was identified as the differential quantity of reads mapped to the rDNA region (See Discussion).

Analysis of the distribution of ATAC-seq reads across the reference genome revealed elevated chromatin accessibility near the transcription start site (TSS) (**Fig. 2b**, top), as reported previously^21^. In spores (0–3h), in addition to the TSS, substantial quantities of ATAC-seq reads were mapped to the coding sequence (CDS) regions of the genes (**Fig. 2b**, top). Their density within the CDS was higher in spores compared to vegetative cells across most genes (**Fig. 2c**, left). Moreover, in germinating spores (1h and 3h), there was an increase in reads on the CDS side compared to dormant spores (**Figs. 2b, c**).

In contrast, rDNA genes exhibited distinct patterns compared to CDS genes. Reflecting the kinetics of the proportion of mapped reads on the rDNA region (**Fig. 2a**), rDNA consistently demonstrated high read density from TSS to the transcription termination site (TTS) during the vegetative phase, yet showed reduced density during the spore phase (**Fig. 2b**, bottom). tRNA genes tended to show similar pattern to rDNA, if not identical (**Fig. 2c**, right).

In the budding yeast *Saccharomyces cerevisiae*, osmotic stress has been shown to trigger transcription of downstream genes by altering nucleosome positioning near promoters^21^. The position and occupancy of nucleosomes can be inferred by classifying reads based on their size in ATAC-seq. To investigate whether nucleosome position sliding occurs in spores compared to vegetative cells, we examined the distribution of reads derived from mono-nucleosomes (**fig. S3**). Our analysis revealed that nucleosome positions remain unchanged regardless of spore germination. Consequently, the increased accessibility observed in the CDS in spores was not associated with a shift in nucleosome positioning. Thus, unlike the example in budding yeast, during germination of *S. pombe*, accessibility changes predominantly occur in the CDS. To our knowledge, this represents the first documented instance of global accessibility changes in the CDS.

### Gene-body peaks vary as germination proceeds

To validate the effect of this increased read density observed on CDS in spores, peak calling analysis was conducted using the software MACS2^26^ (Model-based Analysis of ChIP-Seq 2). Both the narrow and broad options were employed for MACS2: the narrow peak algorithm identifies peaks with a narrow width, facilitating detection of binding sites for proteins such as transcription factors. The broad peak option identifies peaks with greater width than the narrow option, merging adjacent peaks and enabling the detection of peaks associated with histone modifications on gene bodies including CDS. As a result, the narrow option mainly identified peaks upstream of CDS, while the broad option identified peaks spanning CDS (**Fig. 3a**). MACS2 identified approximately 2,400 to 4,800 peaks for each option, with approximately 2,300 to 5,500 of these peaks being common across biological replicates (**Table S3**). In subsequent analyses, peaks shared between the two biological replicates were used (**fig. S4a**). To assess the variation in peak width between these options, the width of each peak was plotted (**Fig. 3b, Table S4**). For spores, the median peak width was found to be between 191 and 195 bp when using the narrow option, and between 354.5 and 382 bp with the broad option. In contrast, for vegetative cells, the median peak width measured 187 bp with the narrow option and 235.5 bp with the broad option, showing that the differences between the two options were smaller than those observed for spores. These results suggest that a key feature of chromatin states in spores may be the presence of broad peaks.

**Figure 3.**
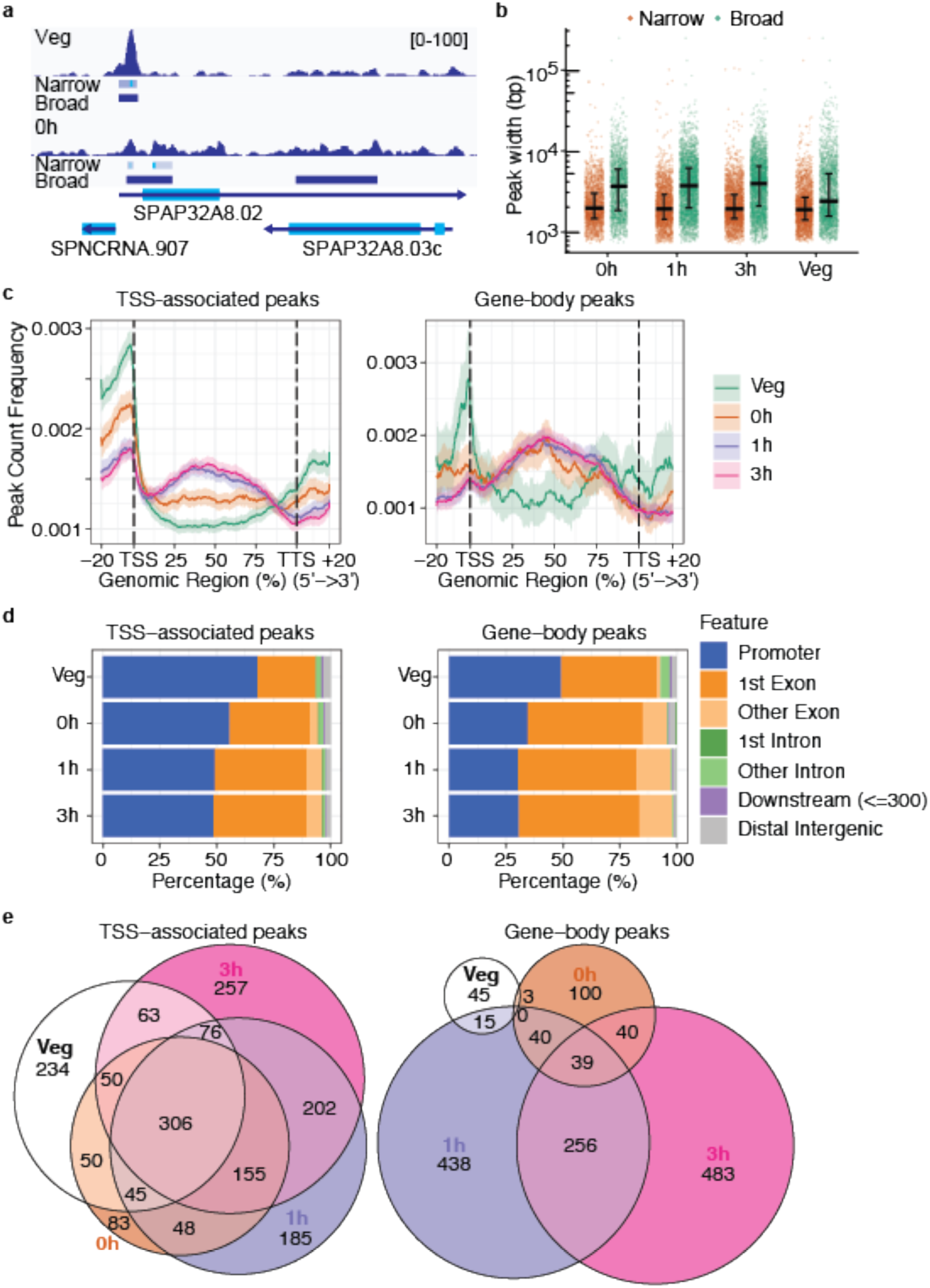
Patterns of ATAC-seq signals in spores after induction of germination reveals broad peaks downstream of the genes. **(a)** An example of the peak range as detected using MACS2. The positions of peaks identified under the two conditions, spore 0h and vegetative cells, are depicted by bars, corresponding to the narrow (pale blue) and broad options (dark blue), respectively. CDS locations and transcriptional directions are shown as in Fig. 2b. **(b)** Each plot depicts the widths of peaks detected under each condition, with median and the range of 25^th^-75^th^ percentiles. Orange, narrow; green, broad. n = 1762-3327 **(c**-**e)** Peaks were categorized into TSS-associated peaks (left) or Gene-body peaks (right) according to the overlaps of narrow and broad option for MACS2 peak calling (see **figs. S4a, b**). **(c)** Average signal intensity of the detected peaks along the gene sequence (TSS–TTS) is presented for each condition. The area corresponding to the 95% confidence interval is indicated by the shaded region. **(d)** Proportion of peaks classified according to the genomic feature. **(e)** Venn diagram showing numbers of genes associated with either peak in the indicated conditions.

To further characterise a feature of peaks specific to spores, peaks detected using narrow and broad options were compared for each condition (**fig. S4b**). All peaks identified using the narrow option were also detected with the broad option. We therefore categorised peaks into two groups: those shared between two options and those uniquely identified by the broad option. The average signal profiles relative to gene body were visualised with these two categories (**Fig.3c**). The peaks commonly identified by both narrow and broad options are located in regions at the transcription start site (TSS) (**Figs. 3a, c**); therefore, these peaks are referred to as “TSS-associated peaks” hereafter. TSS-associated peaks reduced the frequency at TSS on average as germination proceeded (**Fig. 3c**). On the contrary, the peaks uniquely identified by the broad option were found in the gene body including exons (**Figs. 3a, c**). Thus, these peaks are termed “gene-body peaks”. Number of peaks identified exclusively with the broad option significantly increased in germination-induced spores (1h and 3h, **fig. S4b**). More than 1,200 peaks were identified in germination-induced spores, whereas 195 and 341 peaks were identified in vegetative cells and dormant spores, respectively. Considering that the total number of peaks identified in dormant spores (Spore 0h) was the lowest (1,874 peaks), the proportion of peaks was approximately twofold higher in dormant spores than in vegetative cells (9% in veg; 18% in 0h; 38-39% in 1,3h). Consequently, we determined that the gene-body peaks more accurately represent the changes in chromatin states associated with progression of spore germination.

To verify the peak locations, genomic features were assigned to the peaks (**Fig. 3d**). In vegetative cells, TSS-associated peaks detected near the promoter reached highest up to approximately 60%. Instead, in spores, the proportion of gene-body peaks in 1^st^ and other exons reached 70%. This tendency expanded upon germination induction. Consistent with this, peak distances from TSS increased especially in gene-body peaks in germination induced conditions (**fig. S4c**). Thus, differences of vegetative cells and spores regarding ATAC-seq reads are remarkable in the increase of chromatin accessibility particularly in exons.

To characterize genes associated with higher chromatin accessibility during germination, both TSS-associated peaks and gene-body peaks were annotated and mutually compared for overlaps (**Fig. 3e, Table S5**). A distinct pattern upon germination was observed between those two peak types. Within the TSS-associated peaks, each condition exhibited a range of 1021–1440 genes, with 306 genes (17% of the union of genes identified in at least one condition) among them overlapped across all conditions. Only 5-15% of them (83-257 genes) were specifically identified at each condition. In a sharp contrast, among 150–920 genes with gene-body peaks, no gene was commonly detected across all the condition and only 39 genes (3%) were overlapped during spore germination (**Fig. 3e**). Notably, 438 genes (30%) and 483 genes (33%) were exclusively identified in 1h and 3h, respectively. These results suggest that there are two types of peaks; one is TSS-associated peaks that maintain the memory of global gene regulation independently of growth states, the other is gene-body peaks that might be associated with changes in chromatin states during spore germination. The gene-body peaks may be regarded as the most indicative of the variations in chromatin states among vegetative cells and spores.

To further investigate temporal alterations of the chromatin state associated with germination, we conducted a Gene Ontology (GO) analysis to identify the functional genes that constitute TSS-associated or gene-bogy peaks (**Table S6**). For GO terms derived from TSS-associated peaks, 27% of the GO terms (14 terms) detected in vegetative cells were shared among all the conditions and 50% of the GO terms (35 terms) were commonly identified in germination-induced conditions (**fig. S4d**). GO terms commonly shared in all the conditions were related to “cytoplasmic translation”, “canonical glycolysis”, “flocculation”, “regulation of p38MAPK cascade” and “cellular response to osmotic stress”. In contrast to TSS-associated peaks, GO terms identified from gene-body peaks exhibited no overlap among all the conditions (**fig. S4d**). A significant number of GO terms was specific to each condition (72-85%) for gene-body peaks, unlike the case for TSS-associated peaks (25-41%). During germination (1h and 3h), nine GO terms were shared, including “glutamine family amino acid metabolic process” and “regulation of cell growth”. Among the germination stage-specific GO terms, we identified 38 GO terms at 1 h after germination induction, including “urea cycle”, “arginine biosynthetic process via ornithine”, “nucleus organization” and “lipid catabolic process”, and 32 GO terms at 3 h, including “aromatic amino acid family biosynthetic process” and “establishment of bipolar cell polarity”. Thus, chromatin at genes involved in nutrient generation and cell cycle progression may become accessible during germination. This led us to hypothesise that the genes that perform crucial functions vary according to the germination stage, and it is possible that the chromatin state of these genes becomes accessible only at specific stages, thereby facilitating temporal changes in gene expression.

### Cross comparison of RNA-seq and ATAC-seq in GO terms

To test this hypothesis, we performed a comparative analysis of GO term results derived from ATAC-seq data obtained in this study with those from our previous scRNA-seq data^7,8^. Those studies employed scRNA-seq on spores induced to germinate, enabling the deduction of temporal transcriptomic changes during germination: the transcriptional states were classified into seven successive stages (Stages 1 to 7) upon dormancy breaking. To examine temporal transcriptional changes in detail, differentially expressed genes (DEGs) between each pair of consecutive germination stages were identified from our scRNA-seq data, and DEGs were assigned to the stages at which they were highly expressed (**fig. S5a**). We then investigated the overlaps between germination stage-specific DEGs in scRNA-seq and genes associated with ATAC-seq peaks (**Fig.4a**). Genes associated with TSS-associated peaks exhibited a relatively uniform overlap with DEGs throughout germination stages. 19.8-39.4% (18-66 genes) of upregulated DEGs were identified as genes associated with TSS-associated peaks. In contrast, genes associated with gene-body peaks overlapped with DEGs preferentially at specific germination stages. Gene-body peaks identified in 1h and 3h showed preferential overlaps with upregulated DEGs during the later stages of germination (Stages 4-7). These results suggest that genes associated with gene-body peaks correlate with a temporal transcriptional profile during germination.

We further investigated whether changes in chromatin state during germination accompany transcriptional upregulation. At each stage, expression levels of upregulated DEGs were categorized according to chromatin state, based on the TSS-associated or gene-body peaks. When expression data from all single spores were utilized, DEGs associated with TSS-associated peaks showed relatively constant expression, regardless of ATAC-seq peak-detection condition (**fig. S5b**). DEGs associated with gene-body peaks showed similar profiles; however, their expression levels became slightly higher at later germination stages (Stages 4-7) particularly in germination-induced conditions (1h and 3h). We then selected single spores from the stage of interest and neighbouring stages for gene expression quantification (**Fig. 4b**). Even after the selection of stage-specific single spores, expressions of DEGs associated with TSS-associated peaks remained relatively constant during germination. In contrast, DEGs associated with gene-body peaks showed stage-specific increase in expression. At the Stage 1 and 2, DEGs associated with gene-body peaks were highly expressed in dormant spores (0h), whereas the at later stages (Stage 4-7), higher expressions were observed under germination-induced conditions (1h and 3h). These results suggest that DEGs associated with gene-body peaks undergo changes in chromatin state that accompany by their transcriptional upregulations (**Fig. 4c**).

**Figure 4.**
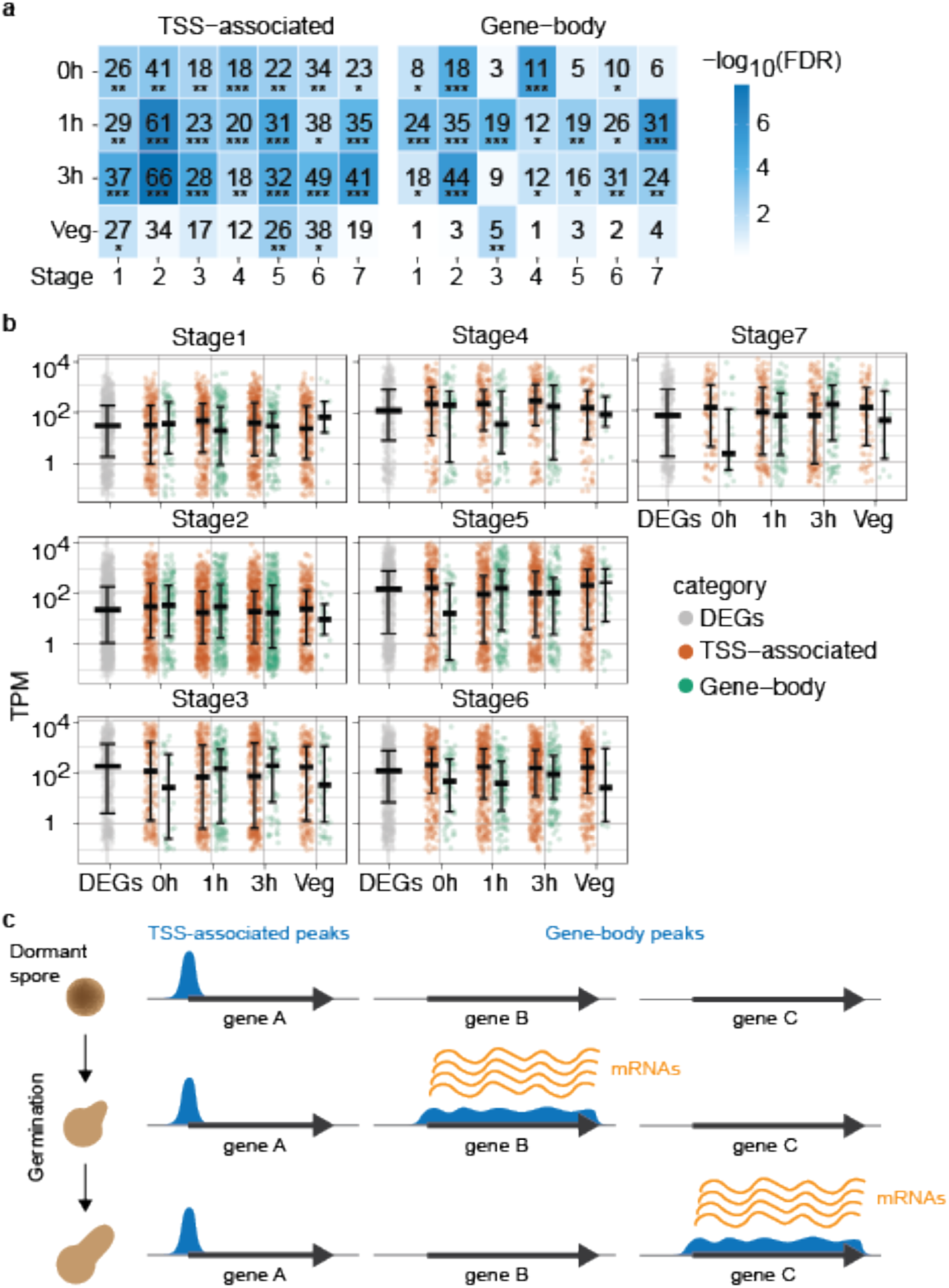
Chromatin dynamics affect gene expression during dormancy breaking. **(a)** Heatmaps showing the numbers of overlapping genes between TSS-associated (left) or gene-body (right) ATAC-seq peaks and differentially expressed genes in scRNA-seq at indicated stages. Colors indicate log_10_ transformed false discovery rate (FDR). ***, FDR < 0.001; **, FDR < 0.01; *, FDR < 0.05. **(b)** Gene expression levels of the DEGs in single spores at indicated stages, with median and the range of 25^th^-75^th^ percentiles. For each stage, at least 23 single spores from the stage of interest and neighbouring were included. Grey, all DEGs; orange, DEGs with TSS-associated peaks; green, DEGs with gene-body peaks. n = 16-6930. **(c)** Genes associated with TSS-associated peaks maintain their open chromatin state throughout the transition from dormancy to germination, whereas those associated with gene-body peaks undergo changes in chromatin accessibility concomitant with their transcriptional activations during germination.

In conclusion, through differential analyses between TSS-associated and gene-body peaks from ATAC-seq, combined with expression profiles from temporal scRNA-seq data, we propose that broadly open chromatin, primarily in the CDS region, may serve as a hallmark indicating transient gene expression during germination.

## Discussion

In this study, we showed that multiple histone components, not only histone H3.c1/Hht1, are transiently reduced during germination of fission yeast spores. Having established an ATAC-seq protocols for spores with *bgs2* mutants, we found that spores displayed a lower fraction of reads mapped to rDNA and tRNA genes and higher accessibility across gene bodies than spheroplasts prepared from vegetative cells. Furthermore, genes covered by gene-body peaks, but not those with TSS-associated peaks, changed as germination progressed and were associated with transcriptional upregulation in a stage-dependent manner in our previously reported scRNA-seq data^7^. These results suggest that gene-body accessibility is a distinctive feature of the chromatin state during spore germination.

In spores, chromatin accessibility was higher across the coding sequence (CDS) of genes following dormancy breaking, whereas nucleosome positions inferred from mono-nucleosome reads remained unchanged (**fig. S3**). This is in contrast to osmotic stress response in budding yeast, in which nucleosome positioning around promoters is altered^21^. Because nucleosome positions did not change, the increased CDS accessibility in spores is therefore not readily explained by nucleosome sliding, for example by chromatin remodelers. Instead, it may reflect changes in nucleosome occupancy or the state of nucleosomes within gene bodies. Histone-GFP signals decreased transiently during the first ∼5 h after the induction of germination by nutritional feeding (**Figs. 1, S1**), a period that encompasses the 1h and 3h time points of our ATAC-seq analysis. This transient reduction may indicate a lower nucleosome density; however, histone-GFP fluorescence cannot distinguish chromatin-bound histones from the soluble histone pool, so we cannot determine whether nucleosome occupancy was actually reduced. Moreover, chemical modifications of histones within nucleosomes may serve as factors influencing alterations in chromatin configuration. For example, the acetylation of lysine residues in the N-terminal region of histone H3 is acknowledged as an indicator of gene expression activation^27,28^. Evaluating the degrees of histone modifications within gene bodies where ATAC-seq signals increased, in dormant and germinating spores over a time course that spans the fluctuation of histone levels, would provide clues to the molecular mechanisms of dormancy breaking.

Mapping ATAC-seq reads to rDNA locus revealed that the highest prevalence of reads in vegetative cells (∼30%) reduced to ∼10% in spores, and tRNA genes showed a similar tendency (**Fig. 2**). These loci are transcribed by different polymerases, RNA polymerase I (rDNA) and III (tRNA genes); therefore, these reductions may have related or independent causes. One possibility is that both reflect reduced growth-related transcription in spores. Actively transcribed rDNA repeats are occupied by more active RNA polymerase I compared to the standard RNA polymerase II, leading to a reduction in nucleosomes^29^, and in proliferating budding yeast cells, ∼60% of total transcription is devoted to rRNA synthesis^30^, which is consistent with the high fraction of ATAC-seq reads at rDNA locus in fission yeast vegetative cells. Assuming that transcriptional activity is reduced in spores relative to vegetative cells, it is plausible that the rDNA region exhibits lower accessibility in spores. Alternatively, considering the variable copy number of repetitive sequences within the rDNA region, there might be a potential reduction in rDNA copy number during the transition from vegetative cells to spore formation via meiosis. If the reduction of accessibility at rDNA reflects reduced transcription, its persistence at 3h, when gene-body accessibility had already changed, raises the possibility that the rDNA and tRNA genes become fully accessible later than the gene-body changes, which can be tested with later time points.

The two classes of peaks showed contrasting behaviors in germinating spores; TSS-associated peaks were relatively stable, while gene-body peaks were highly condition-specific (**Fig. 3**). Of the genes associated with TSS-associated peaks, 306 (17% of union) were common among proliferating vegetative cells, dormant and germinating spores, and the shared GO terms included cytoplasmic translation and glycolysis. Chromatin accessibility at the TSS of these genes may be maintained irrespective of growth state, possibly keeping them poised for rapid use. In contrast, 30% and 33% of genes associated with gene-body peaks were unique to 1h and 3h, respectively. Their GO terms shifted from nutrient metabolism (e.g., urea cycle, arginine biosynthesis) at 1h to establishment of bipolar cell polarity at 3h, in line with an ordered transition from using the supplied nutrients to resuming cell growth.

Consistently, genes associated with gene-body peaks overlapped with upregulated DEGs in a stage-preferential manner and showed stage-dependent increases in expression, whereas genes with TSS-associated peaks overlapped uniformly across stages and showed relatively constant expression (**Fig. 4**). We hypothesized that gene-body peaks in spores marks genes around the time they are upregulated. Although our data do not establish causality or temporal order, the concordance between gene-body accessibility and stage-dependent transcriptional upregulation is consistent with a model in which changes in gene-body chromatin accessibility are coupled to transcriptional activation during germination. Several interpretations remain open: (i) increased accessibility may precede and permit transcription; (ii) transcription itself may increase accessibility within gene bodies; and (iii) both may be driven by a common upstream factor related to germination progression. Because ATAC-seq and scRNA-seq were obtained in separate experiments and compared at the level of time point and stage, our data cannot distinguish among these possibilities. To distinguish these possibilities, ATAC-seq could be performed in the presence of a transcription inhibitor during germination to test whether gene-body accessibility depends on transcription. Moreover, single-cell ATAC-seq combined with single-cell RNA-seq would link accessibility and expression in the same spore.

In summary, broad gene-body accessibility accompanies the transient reduction of histone and stage-dependent gene expression during spore germination, suggesting that chromatin within gene bodies, rather than only at promoters, undergoes changes during the exit from dormancy.

## Acknowledgements

Y.S. was supported by JST SPRING, Grant Number B2R101263201. This work was supported by JSPS KAKENHI Grant Numbers 20K22635 and 22K06092 to Y.K., 21H00261, 23K27173, 25H02582 and 26K02009 to M.S. Support also came from the Dr. Yoshifumi Jigami Memorial Fund to Y.K., Institute for Fermentation, Osaka (IFO) to Y.K. and to M.S., Ohsumi Frontier Science Foundation (OFSF) to M.S. and Waseda University grants for Early Career Researchers Grant Number 2021E-048 to Y.K. and for Special Research Projects Grant Numbers 2020C-738, 2021C-387, 2022C-306, 2023C-283, 2024C-285 to Y.K., 2018B-222, 2019C-570, 2021C-227, 2022C-164, 2023R-026, 2023C-167, 2024C-490, 2025C-154 and 2026C-507 to M.S.

## Author Contribution

Y.S.: Investigation, Methodology, Validation, Visualization, Writing-original draft

H.T.: Investigation

R.A.: Investigation

C.S.: Investigation

M.H.: Resources, Writing-review and editing

H.T.: Writing-review and editing

Y.K.: Conceptualization, Methodology, Data Curation, Project administration, Supervision, Validation, Visualization, Funding Acquisition, Writing-original draft

M.S.: Conceptualization, Funding Acquisition, Project administration, Supervision, Writing-original draft, Writing-review and editing.

## Competing Interests

The authors declare no competing interests.

## Data availability

The sequencing data have been deposited in GEO (accession number GSE355526).

## Materials and Methods

### Yeast strains and protocols

Standard protocols and methodologies were employed for the study of *S. pombe*, as previously documented^31^. The strains used in this study are listed in **Supplemental table 7**. Methods for construction for histone-GFP strains have been illustrated previously^7^. The YE5S medium (yeast extract with supplements: adenine 100 mg/L, uracil 50 mg/L, leucine 50 mg/L, lysine 50 mg/L, histidine 100 mg/L) was used for cultivation of vegetative cells and for refeeding to stimulate spore germination. Sporulation agar plates were used to induce mating, meiosis and sporulation.

### Preparation of spheroplasts

The methods previously described by Flor-Parra et al. (2014)^32^ were modified to prepare spheroplasts of *S. pombe* vegetative cells as follows: cells were cultured in 30 mL YE5S at a concentration of 6‒8 × 10^6^ cells/mL. The cells were subsequently collected and washed twice with 30 mL of SCS buffer. The SCS buffer was prepared by mixing SCSa (20 mM sodium citrate, 1 M D-sorbitol) and SCSb (20 mM citric acid, 1 M D-sorbitol), thereby adjusting pH to 5.8. The cells were then resuspended in 1 mL of SCS containing 0.1 g/mL Lallzyme MMX (Lallmand) and incubated at 30°C for 30 minutes, with stirring every 10 minutes. The cells were then washed once with 1.2 M D-sorbitol and resuspended in 1 mL of 1.2 M D-sorbitol.

### Spore preparation and induction of germination

Prior to initiation of sexual differentiation on a sporulation agar plate, cells were first cultured in YE5S at a density of 0.5-1.0 × 10^7^ cells/mL. 5.0 × 10^7^ cells were harvested, washed four times with sterilized water and resuspended in 50 μL of a Leucine and Uracil mixture (comprising 10 mg/mL Leucine and 1.25 mg/mL Uracil). The cells were subsequently applied to a sporulation agar plate. Following a 3-day incubation at 26.5°C, the resulting spores were used in subsequent experimental procedures. For induction of germination, 2.5 × 10^7^ spores were suspended in 1 mL of 1.2 M D-sorbitol and transferred to either 14 mL of 1.2 M D-sorbitol (designated as 0h) or 14 mL of YE5S medium (designated as 1h or 3h according to the incubation period) at 30°C.

### ATAC-seq library preparation and qPCR quantification

A total of 5 × 10^6^ spores or the same number of spheroplasts prepared from vegetative cells were collected as a pellet in a 1.5 mL tube after washing twice with 1.2 M sorbitol. The cell pellets were suspended by pipetting six times in 50 μL Tn5 reaction mix (Tagmentation buffer (2x) [diagenode] 25 μL, Tagmentase loaded [diagenode] 2.5 μL, 1% digitonin [nacalai tesque] 0.5 μL, 10% Tween 20 [Kanto Chemical] 0.5 μL, 1x PBS (0.23 g/L NaH₂PO₄-2H₂O, 1.16 g/L Na₂HPO₄, 8.5 g/L NaCl) 16.5 μL, H₂O 5 μL). The suspension was then incubated at 37°C for 30 min (Schep et al., 2015). Subsequently, 50 µL of Tn5 stop solution (comprising 1.2 M sorbitol and 50 mM EDTA at pH 8.0) was introduced to terminate the Tn5 reaction.

From the resulting 100 µL suspension, 20 µL was used to determine the numbers of PCR cycles for the indexing PCR (**fig. S2a, Table S1**). To 20 μL of the suspension, 2.5 μL of each primer (Tn5-Fwd and Tn5-Rev, 25 μM) was added, along with 25 μL of NEBNext High-Fidelity 2x PCR Master Mix (NEB). The PCR thermal cycle began with 5 min at 72°C, followed by 30 sec at 98°C, then five cycles of 10 sec at 98°C, 30 sec at 63°C, and 1 min at 72°C, followed by hold at 12 °C. Subsequently, the PCR product was diluted 1,000-fold and 2.5 μL dilution was used as the template for qPCR with primers that included the Tn5 tag sequence but excluded an adapter sequence for sequencing (THUNDER BIRD qPCR mi× 10 μL, Tn5-Fwd 3 μM 2 μL, Tn5-Rev 3 μM 2 μL, 50x ROX reference dye 0.4 μL, H₂O 3.1 μL). The qPCR thermal cycle consisted of an initial step at 95°C for 1 min, followed by 50 cycles of 95°C for 15 sec and 60°C for 1 min. The number of additional cycles was determined based on the fluorescence intensity of amplified products measured at each qPCR cycle. According to the calculation method described in the study by Buenrostro et al. (2015)^33^, the cycles that exceeded one-quarter of the peak fluorescence intensity were counted as additional cycles.

The rest of 80 µL suspension was subjected to purification using the FavorPrep Gel/PCR Purification Mini Kit (Favorgen), and the purified DNA was eluted in 15 µL of elution buffer. To the recovered 15 µL DNA solution, add 5 µL H₂O, 2.5 µL index primer i7, 2.5 µL index primer i5, and 25 µL NEBNext High-Fidelity 2x PCR Master Mix, followed by the execution of PCR. The index primer was prepared using the Nextera® XT Index Kit v2 Set D (96 indexes – 192 samples). The PCR thermal cycle began with 5 min at 72°C, followed by 30 sec at 98°C. This was succeeded by five cycles consisting of 10 sec at 98°C, 30 sec at 63°C, and 1 min at 72°C, before finally cooling to 12°C. During the qPCR process above, the PCR products were maintained on ice. According to the numbers of additional cycles of PCR determined by qPCR (**Table S1**), additional cycles of 10 sec at 98°C, 30 sec at 63°C, and 1 min at 72°C were performed. The resultant sequence library underwent purification using AMPure XP beads. The library preparation, yield, and size selection outcomes were confirmed through TapeStation analysis. Sequencing was executed using the Mi-Seq next-generation sequencer, with paired-end sequencing conducted at 76 base pairs.

### Data processing

Sequenced reads were processed using Trim Galore for filtering and subsequently aligned to the *Schizosaccharomyces pombe* reference genome (ASM294v2.56) using the algorithm bwa mem^34^. Default parameters were applied. PCR duplicates were eliminated using Picard MarkDuplicates. To address the 9-bp offset at the adapter insertion sites caused by the Tn5 transposon, read positions were adjusted using the R/Bioconductor package ATACseqQC^35^. Nucleosome-depleted regions (NDRs) and mononucleosome-associated fragments were characterised by read lengths of 38-100 bp and 140-230 bp, respectively. Unless indicated otherwise (**fig. S3**), only reads corresponding to NDRs were employed for subsequent analyses. Read coverage across biological replicates and aggregate profiles around transcription start sites (TSSs) were generated using deepTools.

To assess the quality of our ATAC-seq data, we reanalyzed previously published ATAC-seq datasets from *S. pombe* (Lei et al. 2021^25^, GSE152086, SRR11963173, 11963174; Schep et al. 2015^21^, GSE66386, SRR1822148, SRR1822149) using the same analytical pipeline as described above.

### Peak calling

Open chromatin regions were identified using MACS2^26^, with and without the application of the --broad option. Peak positions were compared with findOverlapsOfPeaks from ChIPpeakAnno package^36^. Peaks consistently observed across biological replicates were designated as representative peaks for each experimental condition. Representative peak sets obtained with the two MACS2 options were then compared in the same manner. Peak annotation was conducted, average peak profiles relative to gene bodies were generated, and the genomic distribution of peaks and the distance of peaks from transcription start sites (TSSs) were visualized using the R/Bioconductor package ChIPseeker^37^.

### GO term analyses

Gene Ontology (GO) annotation terms were sourced from PomBase. GO enrichment analysis was conducted using the R/Bioconductor package topGO^38^.

### Comparison ATAC peaks with DEGs in scRNA-seq

Differentially expressed genes (DEGs) werer obtained from Tsuyuzaki et al., 2020^7^, in which individual spores are ordered along a virtual time line and assigned to seven germination stages. For each sample or developmental stage, the statistical significance of the overlap between ATAC-seq peak-associated genes and DEGs was evaluated using one-sided Fisher’s exact tests. Obtained *P* values were adjusted for multiple testing using the Benjamini–Hochberg procedure, and adjusted P values (FDR) < 0.05 were considered statistically significant. To visualize germination stage-specific gene expressions, at least 16 single spores from the stage of interest and neighbouring stages were pooled (n = 16-23 spores per group). The expressions of the selected DEGs are displayed for genes associated with TSS-associated or gene-body peaks.

### Fluorescence microscopy

Cells expressing Hht1(H3.c1)-GFP, Hta1(H2A)-GFP, Htb1(H2B)-GFP, and Hhf1(H4.c1)-GFP were placed on SPA plates to induce sporulation. The resultant spore cells were suspended in 200 µL of ddH_2_O, and 7.5-µL aliquots were mounted on multi-well 35-mm glass-bottomed dishes that had been pre-coated with soybean lectin. The wells were then positioned on the stage of a DeltaVision microscope system, as previously described^39^. Following the acquisition of images for the first time point, the wells were replenished with 500 µL of the nutrient-rich medium YE5S. Subsequently, the cells were refocused, and time-lapse observations were conducted at 5-min intervals over a 12-h period.

### Fluorescence measurement

Cells expressing Hht1(H3.c1)-GFP (n=15 cells), Hta1(H2A)-GFP (n=14), Htb1(H2B)-GFP (n=15), and Hhf1(H4.c1)-GFP (n=18) were used for quantification. Image processing and analysis were performed following a sequential workflow using Fiji^40^. Initially, a sum stack was generated from the R3D_D3D.DV file. The R3D_REF.dv file was then converted from 16-bit to 32-bit format. Subsequently, channels were merged, and spores were selected, with the resulting ROI set saved as RoiSet.zip. The merged image was split to isolate the GFP channel, followed by background subtraction using a rolling ball radius of 10.0 pixels. A montage was created by arranging all time-lapse images into a single column in a row-wise manner. Thresholding was applied using the default settings, and the image was converted to a binary mask. Particle analysis was conducted to identify nuclei, and the generated masks were transferred to the montage. Unwanted particles were removed before measuring intensities, areas, and other relevant parameters.

## Supplemental figures

**Supplemental figure 1.**
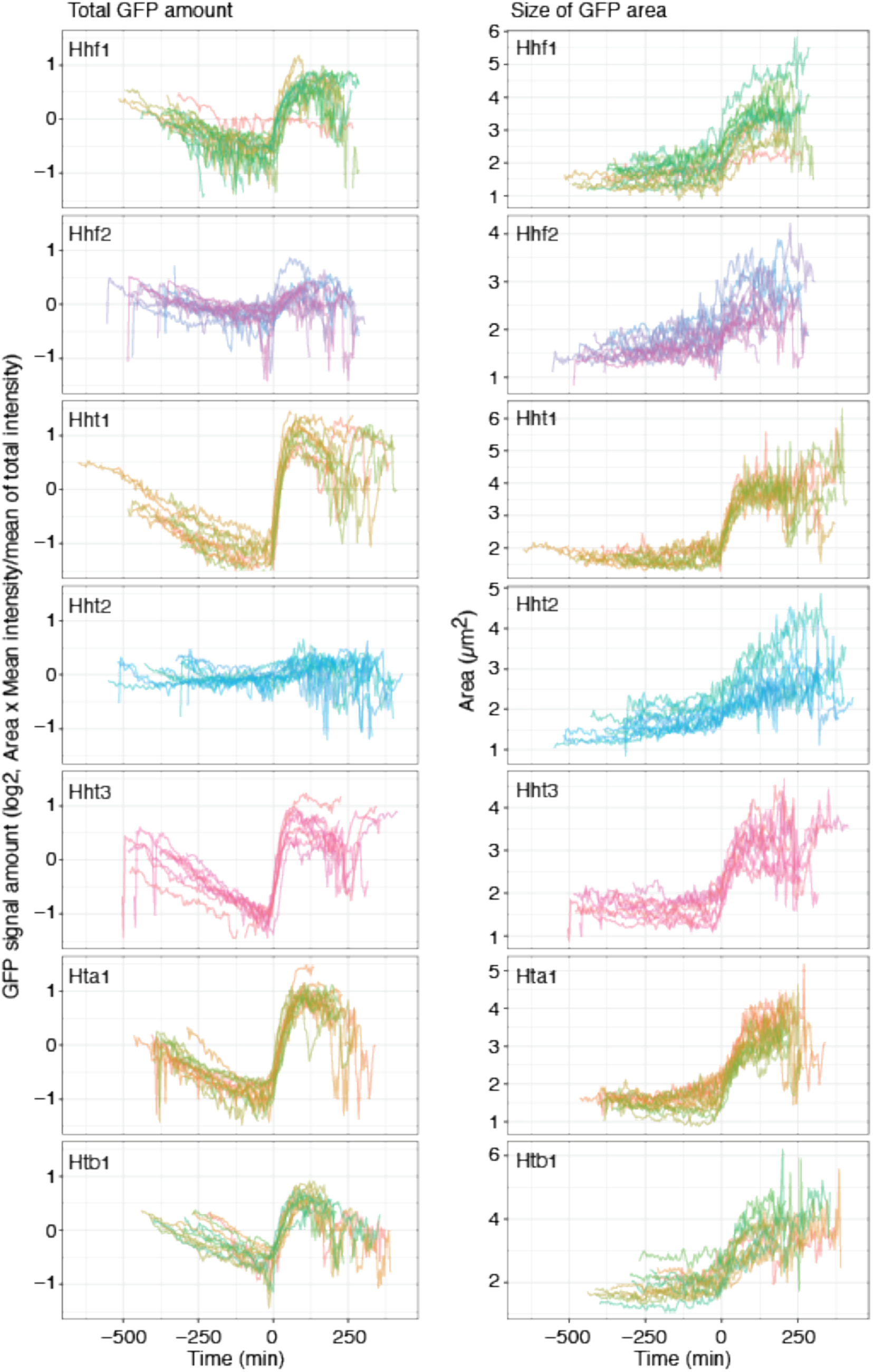
The majority of histone-GFP exhibits temporal fluctuations in fluorescence intensity during the germination process, with corresponding changes in the area of detectable GFP. Time-lapse observations of germination in strains expressing histone–GFP for indicated histone genes was conducted and the fluorescence intensities and their detectable area are monitored over time. n = 15 cells (Hht1-GFP), 14 (Hht2-GFP), 11 (Hht3-GFP), 18 (Hhf1-GFP), 17 (Hhf2-GFP), 15 (Hta1-GFP) and 14 (Htb1-GFP). Time 0 (min) is designated as the point at which a discernible change in brightness was observed.

**Supplemental figure 2.**
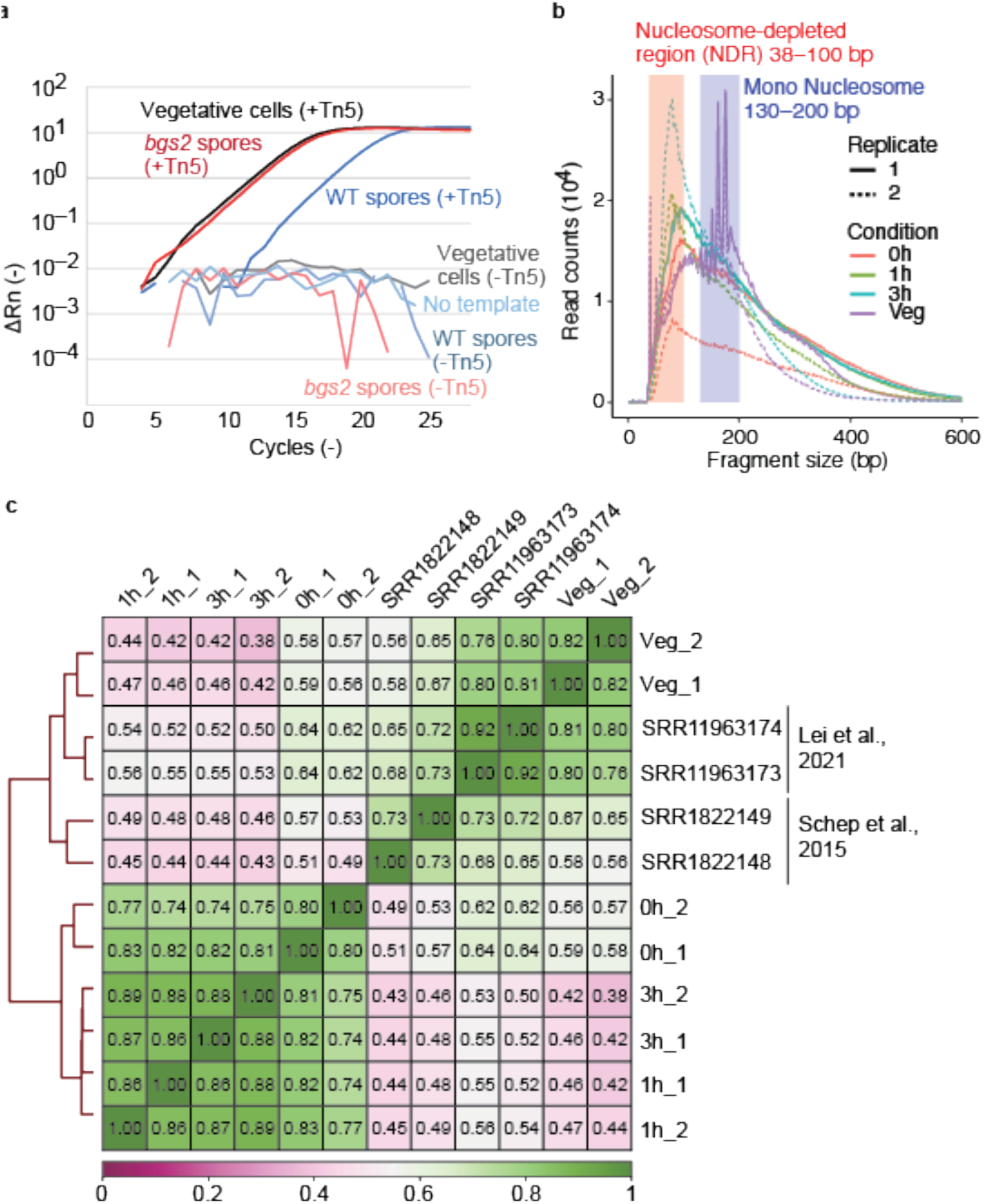
Quality control of ATAC-seq libraries. **(a)** qPCR Verification of Tn5 tagmentation in spores using the methods previously employed with vegetative cells^21,33^. Spheroplasts prepared from vegetative cells and spores of wild type (WT) and *bgs2* mutant were incubated in Tn5 reaction mix with (+Tn5) or without (–Tn5) Tagmentase, followed by five cycles of PCR amplification. Tagmented fragments were quantified by qPCR with Tn5 tag-specific adapter-free primers. **(b)** The distribution of fragment sizes for ATAC-seq samples. Fragments ranging from 38 to 100 bp are presumed to originate from nucleosome-free regions, whereas fragments within the 130 to 200 bp range are considered to originate from mono-nucleosomes. **(c)** The correlation coefficient was calculated for the number of the reads ranging from 38 to 100 bp and mapped to 1 kb bins under the Spearman correlation method. Data from previous studies were incorporated [Schep et al., 2015^21^, SRR1822148 and SRR1822149; Lei et al., 2021^25^, SRR11963173 and SRR11963174]. The intensity of the color within the box indicates the magnitude of the correlation coefficient. The final digit indicates the biological replicate number.

**Supplemental figure 3.**
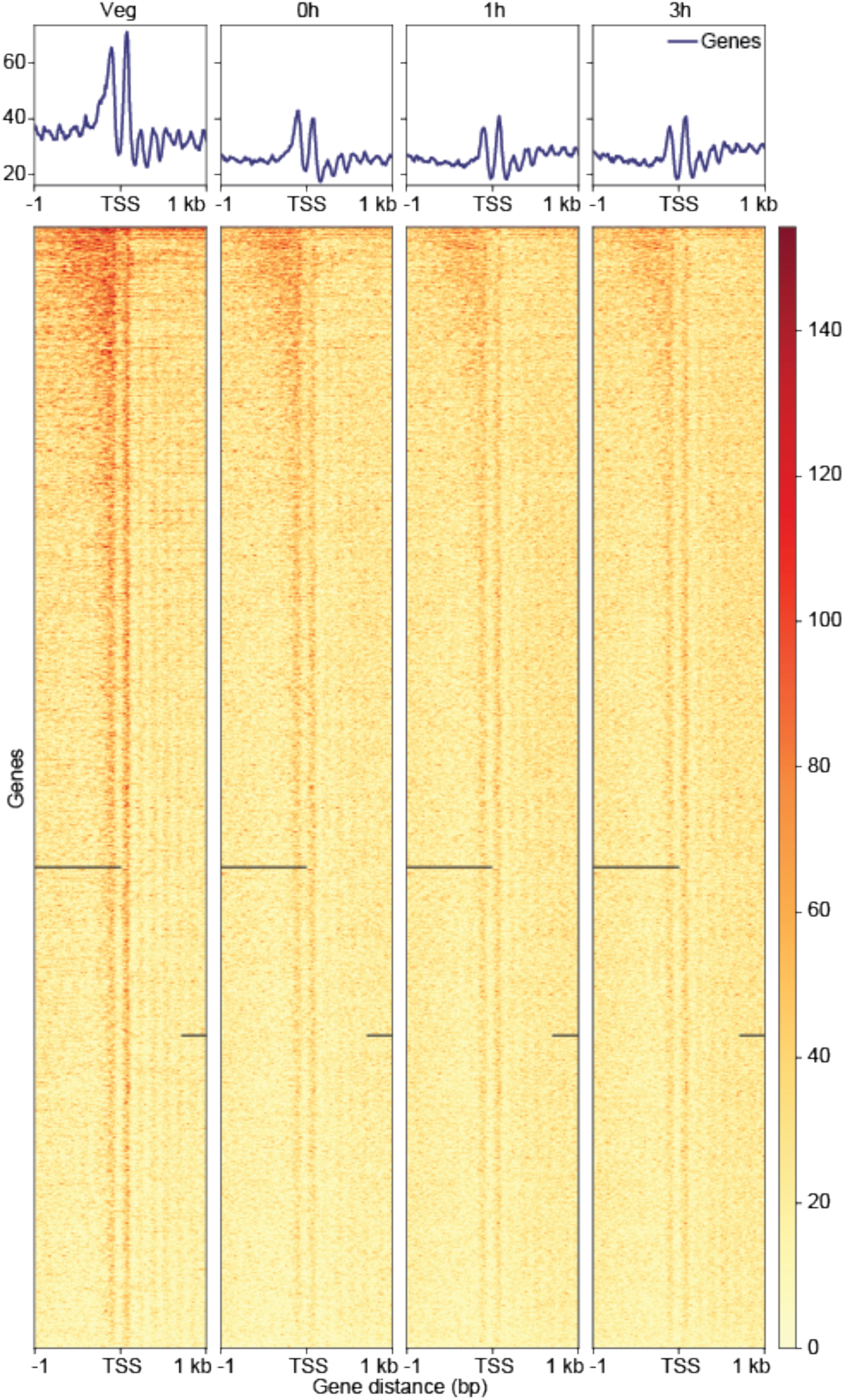
No alterations were detected in the nucleosome positioning within the CDS regions of genes. The distribution of reads corresponding to mono-nucleosomes (130-200 bp) per gene was arranged vertically around the transcription start site (TSS ± 1.0 kb).

**Supplemental figure 4.**
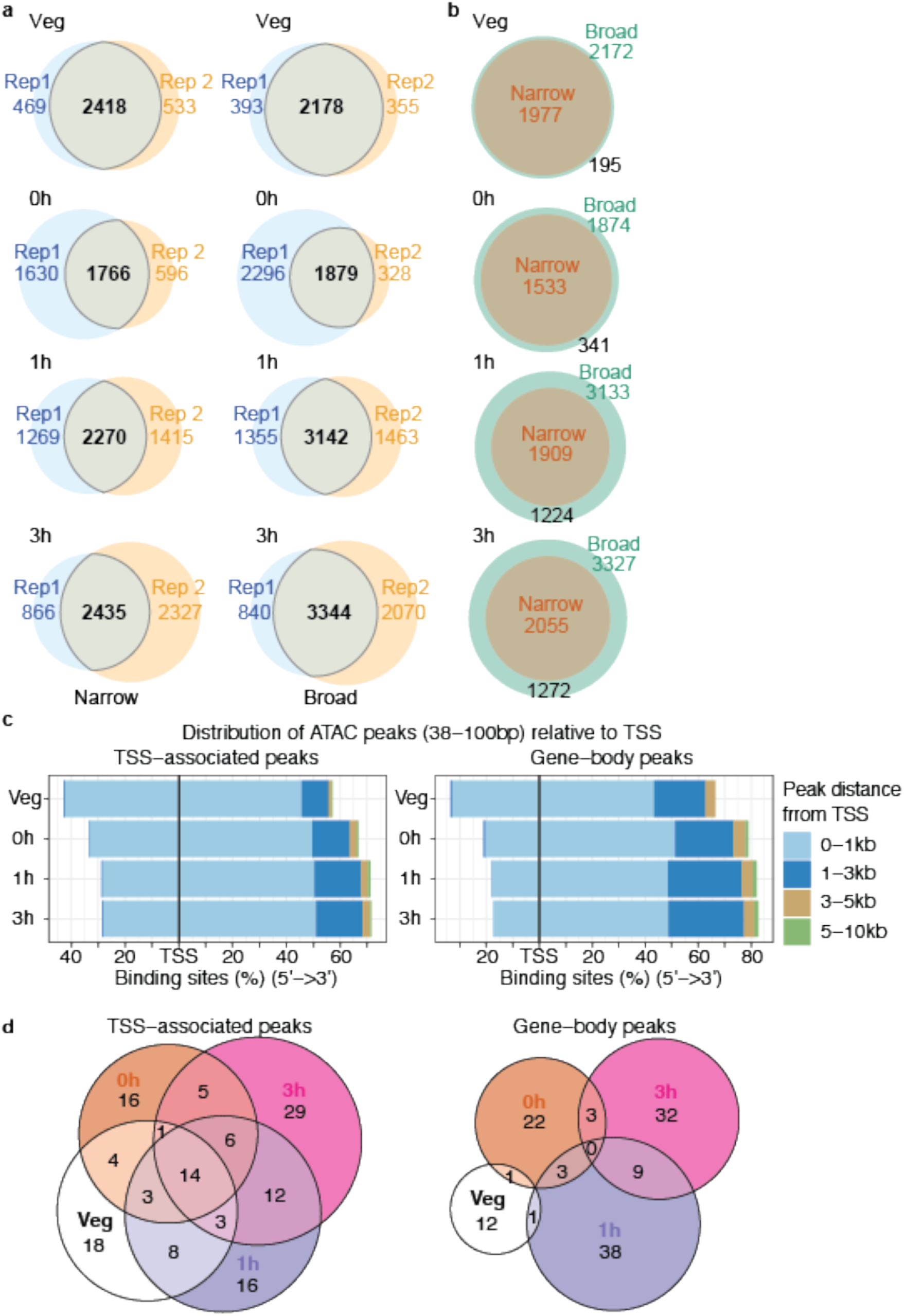
Classifications of peaks determined by MACS2 narrow and broad options and peak locations from TSS. **(a)** Venn diagrams illustrate the overlap between replicates for peaks. For peak calling by MACS2, both narrow (left) and broad (right) options were employed. The number of peaks in specific to each replicate is represented by the corresponding color, while the black values indicate the overlaps between the two biological replicates. These overlapping peaks were used as peaks in each condition for the following analyses. **(b)** Venn diagrams illustrating the connotation of the narrow versus broad options for MACS2 peak calling. The number of peaks in each peak detection option is represented by the corresponding color, while the black values indicate the numbers of peaks detected only in broad option. **(c)** The percentages of peaks classified by their distance from the TSS are shown. Of the two termini representing the region of a peak, the terminus proximal to the TSS was selected for classification according to the distance from the TSS site. **(d)** Venn diagram showing GO terms overlapped among the conditions, separated by genes associated with TSS-associated (left) and gene-body peaks (right).

**Supplemental Figure 5.**
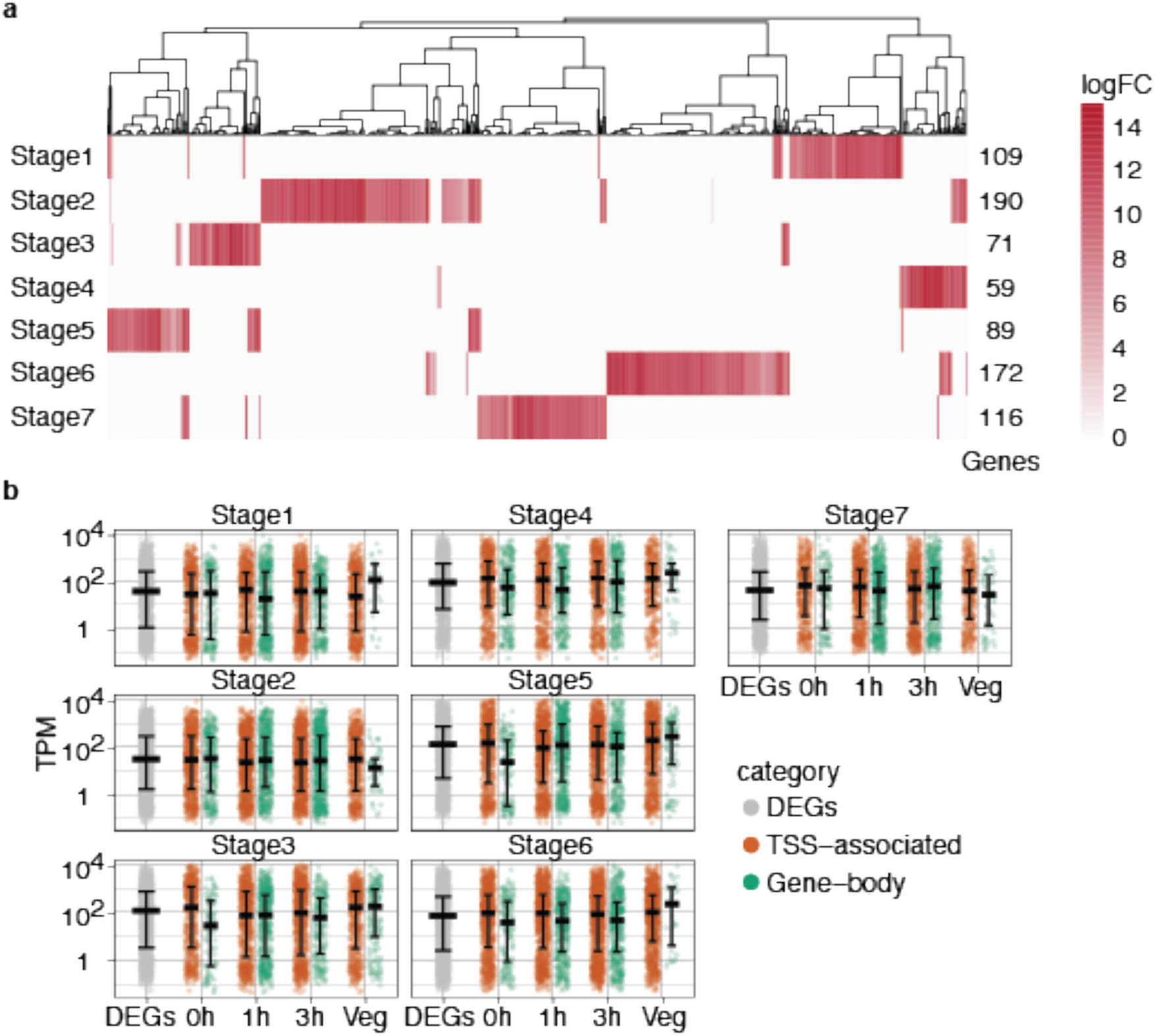
Gene expression profiles during germination determined by scRNA-seq. **(a)** Gene expression profiles categorized by seven virtual stages in scRNA-seq^7,8^. Highly expressed genes were identified by comparing gene expressions at two consecutive stages. **(b)** Gene expression levels of the DEGs shown in fig. S5a across all single spores, with median and the range of 25^th^-75^th^ percentiles. Grey, all DEGs; orange, DEGs with TSS-associated peaks; green, DEGs with gene-body peaks. n = 1140-49560.

---

**Supplemental Table 1.** Summary of PCR cycles used for indexing and ATAC-seq reads.

| Sample ID | Condition | Replicate | Number of additional cycles | Read pairs examined | Unmapped reads | Read pair duplicates | Duplication rate | Estimated library size |
|---|---|---|---|---|---|---|---|---|
| 164 | Vegetative | 1 | 7 | 6,317,247 | 226,394 | 3,559,979 | 0.56 | 3,202,936 |
| 169 | Spore 0h | 1 | 10 | 6,634,851 | 280,005 | 2,961,373 | 0.45 | 4,999,896 |
| 171 | Spore 1h | 1 | 8 | 5,436,166 | 34,948 | 1,635,556 | 0.30 | 7,113,857 |
| 179 | Spore 3h | 1 | 7 | 5,106,414 | 59,838 | 3,403,118 | 0.67 | 1,811,386 |
| 162 | Vegetative | 2 | 10 | 5,568,549 | 36,470 | 1,544,484 | 0.28 | 8,082,261 |
| 170 | Spore 0h | 2 | 13 | 5,994,227 | 43,779 | 2,724,687 | 0.46 | 4,390,661 |
| 172 | Spore 1h | 2 | 11 | 5,683,486 | 35,985 | 1,610,781 | 0.28 | 8,027,215 |
| 180 | Spore 3h | 2 | 9 | 6,074,377 | 44,169 | 2,389,208 | 0.39 | 5,526,397 |

**Supplemental Table 2.**
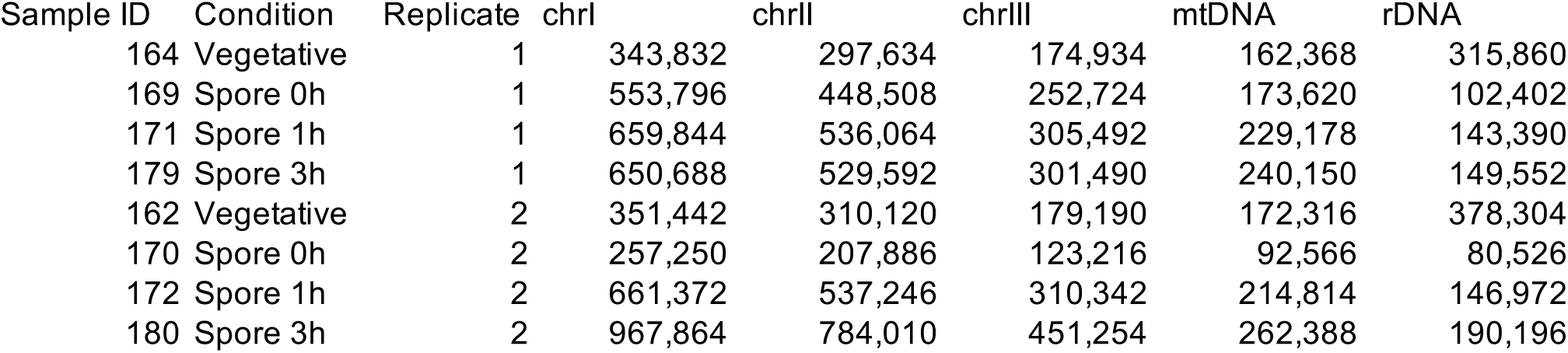
Numbers of reads mapped to chromosomes, mitochondrial DNA and rDNA regions.

**Supplemental Table 3.**
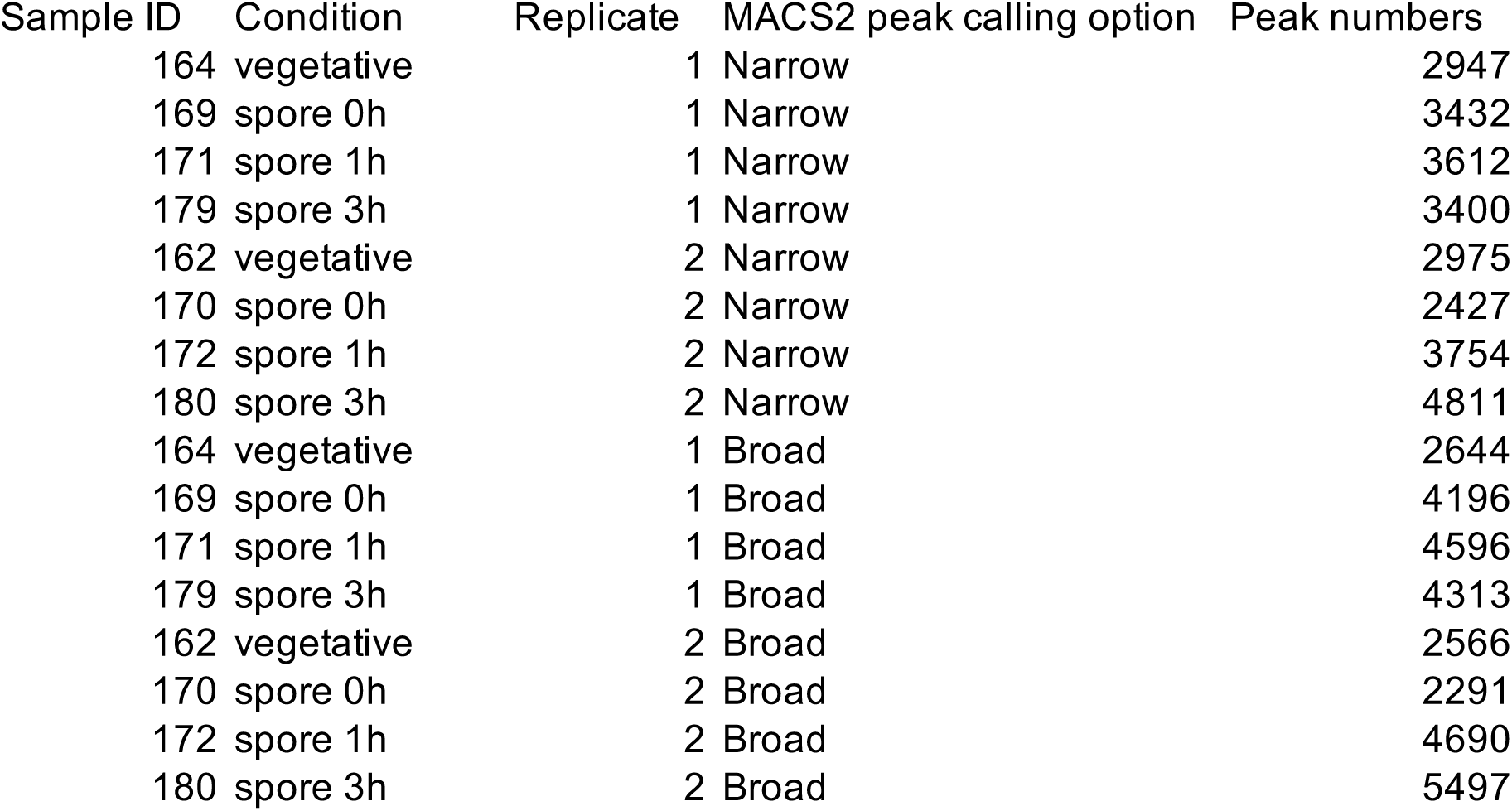
Numbers of peaks identified using MACS2 narrow- and broad-options.

**Supplemental Table 4.** Distribution of peak widths.

| Condition | MACS2 peak calling option | Min width (bp) | 25th percentile width (bp) | Median width (bp) | 75th percentile width (bp) | Max width (bp) |
|---|---|---|---|---|---|---|
| vegetative | Narrow | 77 | 141 | 187 | 262 | 24554 |
| spore 0h | Narrow | 75 | 147 | 195 | 293 | 6830 |
| spore 1h | Narrow | 75 | 144 | 191 | 283.5 | 10380 |
| spore 3h | Narrow | 74 | 147 | 191 | 282 | 10784 |
| vegetative | Broad | 75 | 156 | 235.5 | 500 | 24555 |
| spore 0h | Broad | 76 | 183 | 354.5 | 566.75 | 24546 |
| spore 1h | Broad | 76 | 196 | 360 | 583 | 24547 |
| spore 3h | Broad | 74 | 206 | 382 | 615 | 24552 |

**Supplemental Table 5.** Numbers of annotations associated with ATAC-seq peaks.

| Sample | Peak category | Protein_coding | ncRNA | tRNA | rRNA | snoRNA | snRNA |
|---|---|---|---|---|---|---|---|
| vegetative | TSS-associated | 1402 | 1061 | 95 | 21 | 10 | 5 |
| Spore0h | TSS-associated | 1021 | 769 | 28 | 12 | 1 | 1 |
| Spore1h | TSS-associated | 1296 | 862 | 16 | 6 | 3 | 1 |
| Spore3h | TSS-associated | 1440 | 878 | 19 | 4 | 2 | 0 |
| vegetative | Gene-body | 150 | 107 | 7 | 0 | 0 | 0 |
| Spore0h | Gene-body | 252 | 136 | 6 | 3 | 2 | 0 |
| Spore1h | Gene-body | 888 | 435 | 3 | 2 | 0 | 1 |
| Spore3h | Gene-body | 920 | 479 | 4 | 1 | 0 | 0 |

**Supplemental Table 6.**
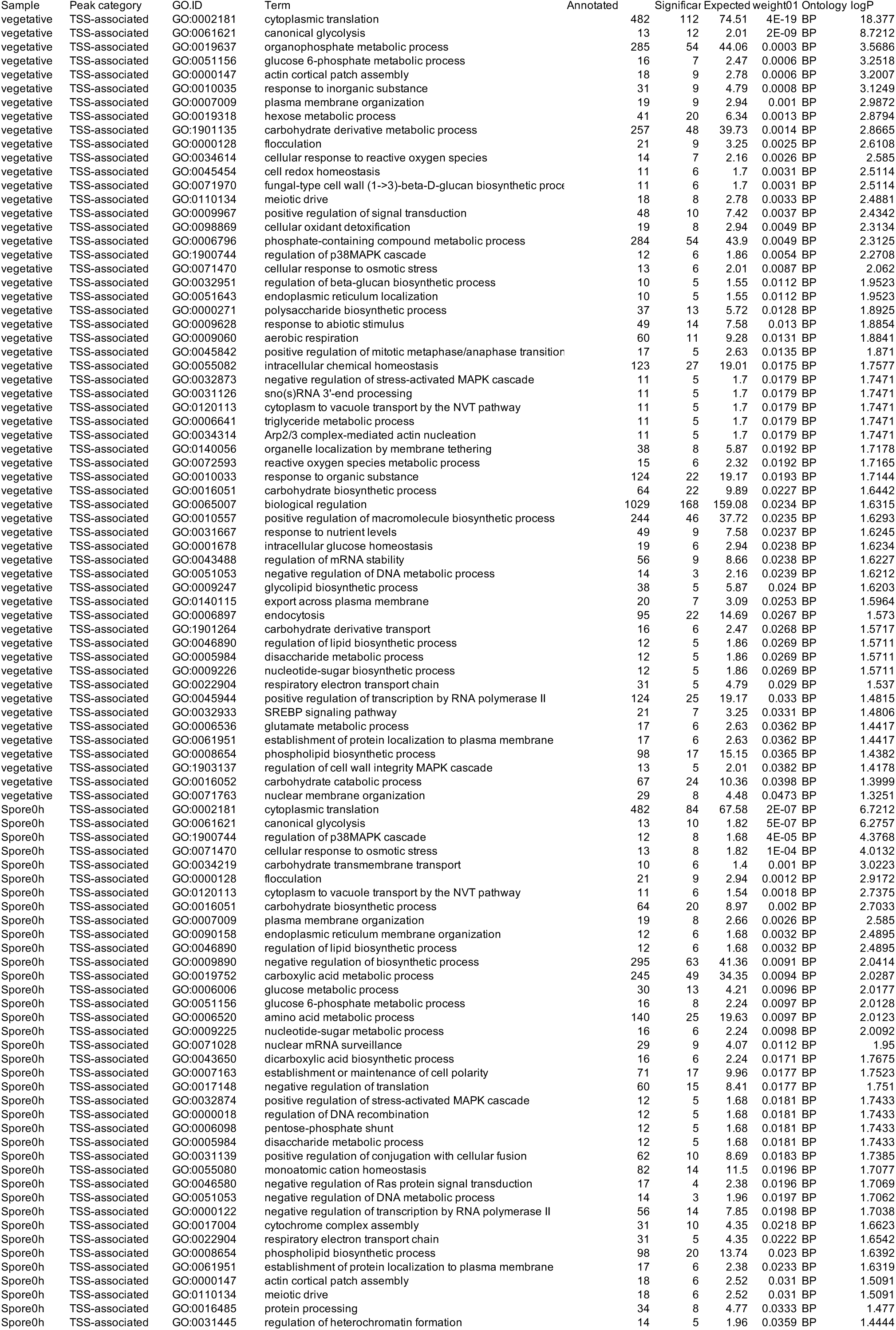
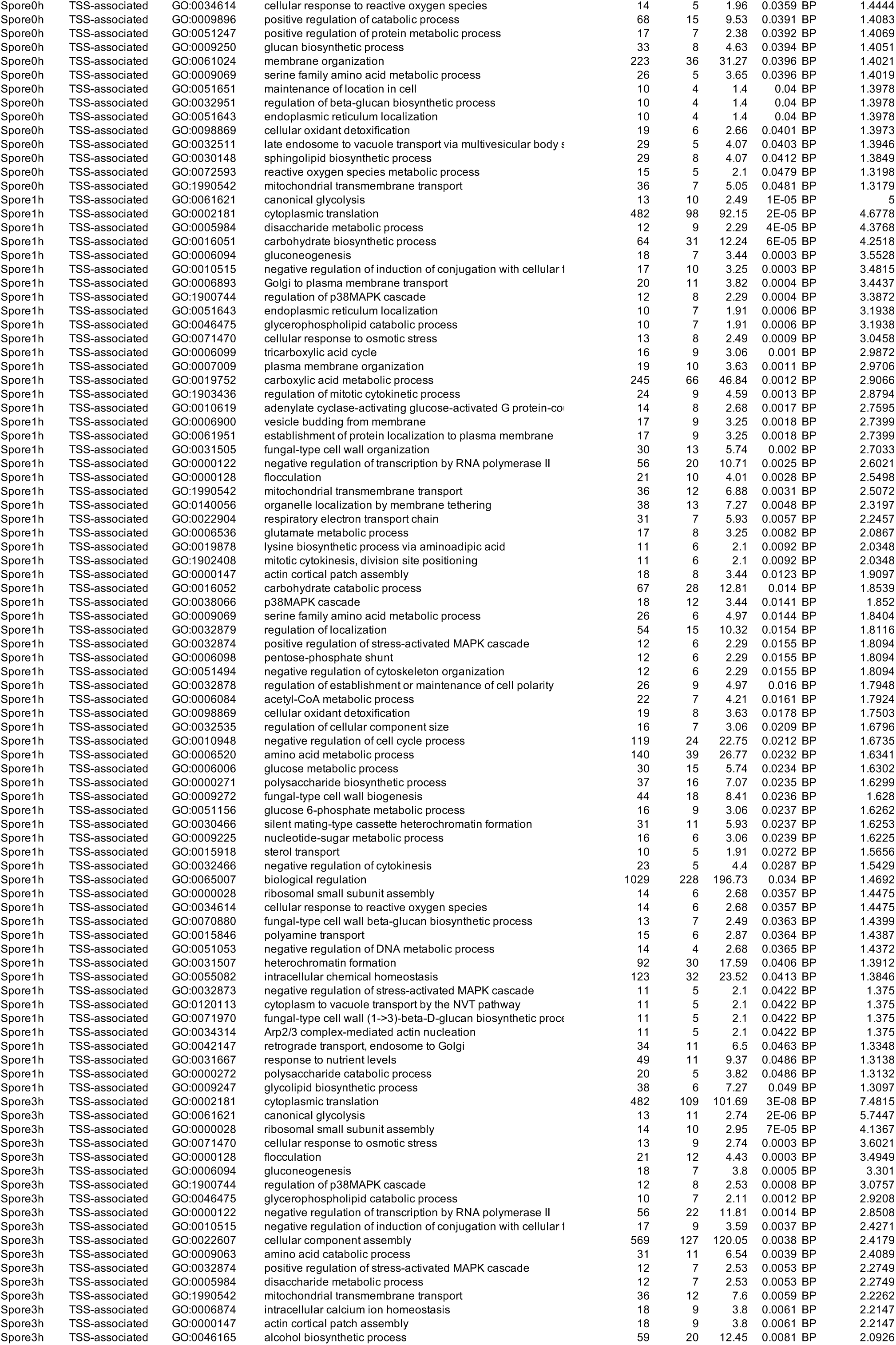
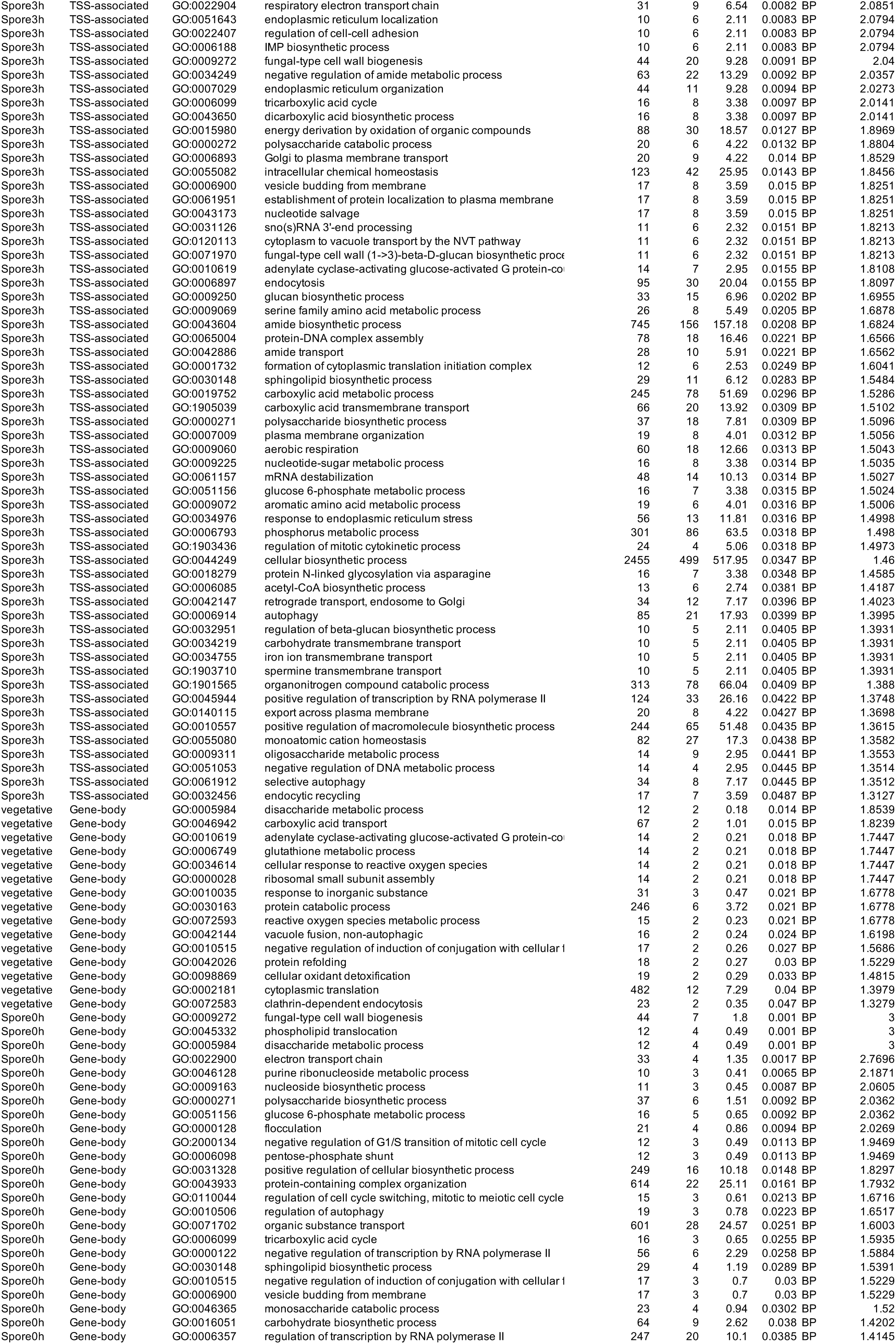
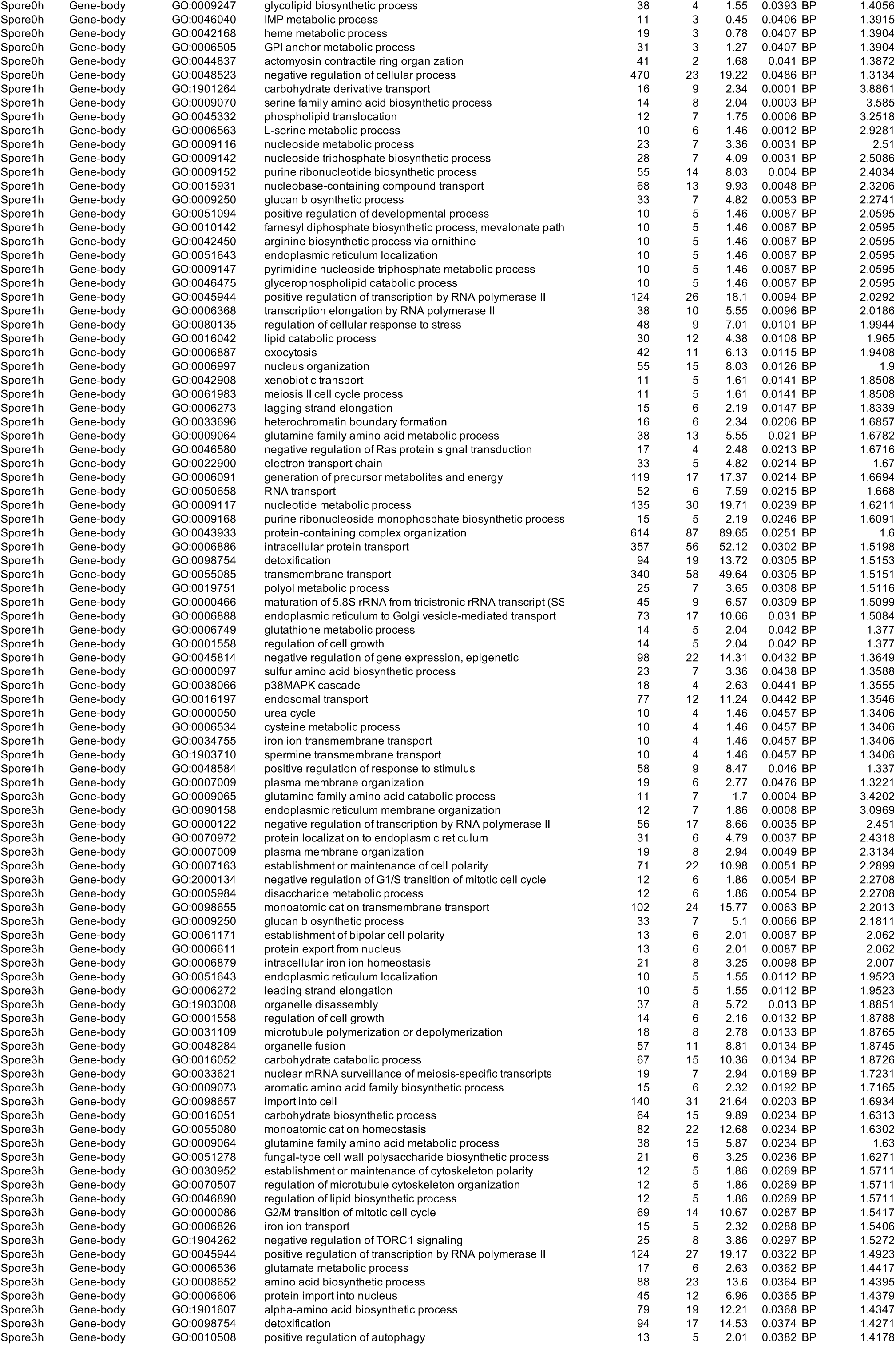
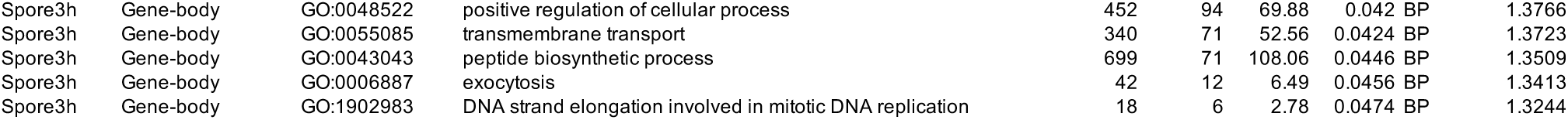
GO terms identified under the indicated conditions.

**Supplemental Table 7.**
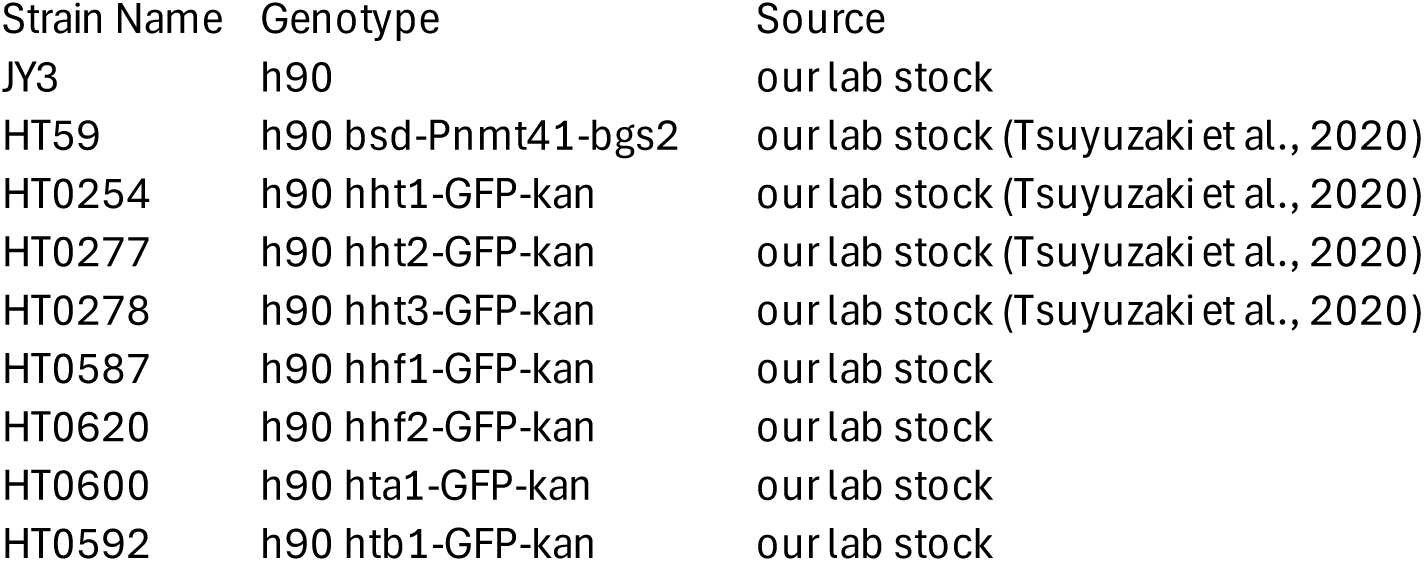
List of fission yeast strains used in this study.

## Notes

### Competing Interest Statement

The authors have declared no competing interest.

## References

1. Rosswag de Souza, S., Böke, E. & Zaffagnini, G. Proteostasis in cellular dormancy: lessons from yeast to oocytes. Trends Biochem. Sci. 50, 646–662 (2025).

2. Su, S. S. Y., Tanaka, Y., Samejima, I., Tanaka, K. & Yanagida, M. A nitrogen starvation-induced dormant G0 state in fission yeast: the establishment from uncommitted G1 state and its delay for return to proliferation. J. Cell Sci. 109, 1347–1357 (1996).

3. Sun, S. & Gresham, D. Cellular quiescence in budding yeast. Yeast 38, 12–29 (2021).

4. Ohtsuka, H., Imada, K., Shimasaki, T. & Aiba, H. Sporulation: A response to starvation in the fission yeast *Schizosaccharomyces pombe*. MicrobiologyOpen 11, e1303 (2022).

5. Hatanaka, M. & Shimoda, C. The cyclic AMP/PKA signal pathway is required for initiation of spore germination in *Schizosaccharomyces pombe*. Yeast 18, 207–217 (2001).

6. Sakai, K., Kondo, Y., Aoki, K. & Goto, Y. Molecular and biophysical perspectives on dormancy breaking: lessons from yeast spore. Biomolecules 15, 701 (2025).

7. Tsuyuzaki, H. et al. Time-lapse single-cell transcriptomics reveals modulation of histone H3 for dormancy breaking in fission yeast. Nat. Commun. 11, 1265 (2020).

8. Tsuyuzaki, H., Ujiie, R. & Sato, M. Wake-up alarm: virtual time-lapse gene expression landscape illuminates mechanisms underlying dormancy breaking of germinating spores. Curr. Genet. 67, 519–534 (2021).

9. Swygert, S. G. et al. Condensin-dependent chromatin compaction represses transcription globally during quiescence. Mol. Cell 73, 533–546 (2019).

10. Swygert, S. G. et al. Local chromatin fiber folding represses transcription and loop extrusion in quiescent cells. eLife 10, e72062 (2021).

11. Tan, Z. Y. et al. Macromolecular and cytological changes in fission yeast G0 nuclei. J. Cell Sci. 138, jcs263654 (2025).

12. Kornberg, R. D. Chromatin structure: a repeating unit of histones and DNA. Science 184, 868–871 (1974).

13. Kornberg, R. D. & Lorch, Y. Chromatin structure and transcription. Annu. Rev. Cell Biol. 8, 563–587 (1992).

14. Boyle, A. P. et al. High-resolution mapping and characterization of open chromatin across the genome. Cell 132, 311–322 (2008).

15. Sabo, P. J., et al. Genome-wide identification of DNaseI hypersensitive sites using active chromatin sequence libraries. Proc. Natl Acad. Sci. USA 101, 4537–4542 (2004).

16. Sabo, P. J. et al. Genome-scale mapping of DNase I sensitivity in vivo using tiling DNA microarrays. Nat. Methods 3, 511–518 (2006).

17. Giresi, P. G., Kim, J., McDaniell, R. M., Iyer, V. R. & Lieb, J. D. FAIRE (formaldehyde-assisted isolation of regulatory elements) isolates active regulatory elements from human chromatin. Genome Res. 17, 877–885 (2007).

18. Schones, D. E. et al. Dynamic regulation of nucleosome positioning in the human genome. Cell 132, 887–898 (2008).

19. Song, L. & Crawford, G. E. DNase-seq: a high-resolution technique for mapping active gene regulatory elements across the genome from mammalian cells. Cold Spring Harb. Protoc. 2010, pdb.prot5384 (2010).

20. Buenrostro, J. D., Giresi, P. G., Zaba, L. C., Chang, H. Y. & Greenleaf, W. J. Transposition of native chromatin for fast and sensitive epigenomic profiling of open chromatin, DNA-binding proteins and nucleosome position. Nat. Methods 10, 1213–1218 (2013).

21. Schep, A. N. et al. Structured nucleosome fingerprints enable high-resolution mapping of chromatin architecture within regulatory regions. Genome Res. 25, 1757–1770 (2015).

22. Corces, M. R. et al. An improved ATAC-seq protocol reduces background and enables interrogation of frozen tissues. Nat. Methods 14, 959–962 (2017).

23. Liu, J., Tang, X., Wang, H. & Balasubramanian, M. Bgs2p, a 1,3-β-glucan synthase subunit, is essential for maturation of ascospore wall in *Schizosaccharomyces pombe*. FEBS Lett. 478, 105–108 (2000).

24. Martín, V., Ribas, J. C., Carnero, E., Durán, A. & Sánchez, Y. *bgs2*+, a sporulation-specific glucan synthase homologue is required for proper ascospore wall maturation in fission yeast. Mol. Microbiol. 38, 308–321 (2000).

25. Lei, B. et al. A synthetic approach to reconstruct the evolutionary and functional innovations of the plant histone variant H2A.W. Curr. Biol. 31, 182–191 (2021).

26. Zhang, Y. et al. Model-based analysis of ChIP-Seq (MACS). Genome Biol. 9, R137 (2008).

27. Kouzarides, T. Chromatin modifications and their function. Cell 128, 693–705 (2007).

28. Tan, M. et al. Identification of 67 histone marks and histone lysine crotonylation as a new type of histone modification. Cell 146, 1016–1028 (2011).

29. Scheer, U. Changes of nucleosome frequency in nucleolar and non-nucleolar chromatin as a function of transcription: an electron microscopic study. Cell 13, 535–549 (1978).

30. Warner, J. R. The economics of ribosome biosynthesis in yeast. Trends Biochem. Sci. 24, 437–440 (1999).

31. Moreno, S., Klar, A. & Nurse, P. Molecular genetic analysis of fission yeast *Schizosaccharomyces pombe*. Methods Enzymol. 194, 795–823 (1991).

32. Flor-Parra, I., Zhurinsky, J., Bernal, M., Gallardo, P. & Daga, R. R. A Lallzyme MMX-based rapid method for fission yeast protoplast preparation. Yeast 31, 61–66 (2014).

33. Buenrostro, J. D., Wu, B., Chang, H. Y. & Greenleaf, W. J. ATAC-seq: a method for assaying chromatin accessibility genome-wide. Curr. Protoc. Mol. Biol. 109, 21.29.1–21.29.9 (2015).

34. Li, H. & Durbin, R. Fast and accurate short read alignment with Burrows–Wheeler transform. Bioinformatics 25, 1754–1760 (2009).

35. Ou, J. et al. ATACseqQC: a Bioconductor package for post-alignment quality assessment of ATAC-seq data. BMC Genomics 19, 169 (2018).

36. Zhu, L. J. et al. ChIPpeakAnno: a Bioconductor package to annotate ChIP-seq and ChIP-chip data. BMC Bioinformatics 11, 237 (2010).

37. Yu, G., Wang, L.-G. & He, Q.-Y. ChIPseeker: an R/Bioconductor package for ChIP peak annotation, comparison and visualization. Bioinformatics 31, 2382–2383 (2015).

38. Alexa, A., Rahnenführer, J. & Lengauer, T. Improved scoring of functional groups from gene expression data by decorrelating GO graph structure. Bioinformatics 22, 1600–1607 (2006).

39. Sato, M., Toya, M. & Toda, T. Visualization of fluorescence-tagged proteins in fission yeast: the analysis of mitotic spindle dynamics using GFP-tubulin under the native promoter. Methods Mol. Biol. 545, 185–203 (2009).

40. Schindelin, J. et al. Fiji: an open-source platform for biological-image analysis. Nat. Methods 9, 676–682 (2012).

